# A generalizable deep learning framework for inferring fine-scale germline mutation rate maps

**DOI:** 10.1101/2021.10.25.465689

**Authors:** Yiyuan Fang, Shuyi Deng, Cai Li

**Affiliations:** State Key Laboratory of Biocontrol, School of Life Sciences, Sun Yat-sen University, Guangzhou, Guangdong, China

## Abstract

Germline mutation rates are essential for genetic and evolutionary analyses. Yet, estimating accurate fine-scale mutation rates across the genome is a great challenge, due to relatively few observed mutations and intricate relationships between predictors and mutation rates. Here we present MuRaL (<u>Mu</u>tation <u>Ra</u>te <u>L</u>earner), a deep learning framework to predict mutation rates at the nucleotide level using only genomic sequences as input. Harnessing human germline variants for comprehensive assessment, we show that MuRaL achieves better predictive performance than current state-of-the-art methods. Moreover, MuRaL can build models with relatively few training mutations and a moderate number of sequenced individuals, and can leverage transfer learning to further reduce data and time demands. We apply MuRaL to produce genome-wide mutation rate maps for four representative species - *Homo sapiens, Macaca mulatta, Arabidopsis thaliana* and *Drosophila melanogaster*, demonstrating its high applicability. As an example, we use improved mutation rate estimates to stratify human genes into distinct groups which are enriched for different functions, and highlight that many developmental genes are subject to high mutational burden. The open-source software and generated mutation rate maps can greatly facilitate related research.

## Introduction

Germline *de novo* mutations (DNMs), which occur either during gametogenesis or post-zygotically, are crucial for evolution and play important roles in many human diseases^1^. Average *de novo* mutation rates for single nucleotide variants (SNVs) in the human genome were estimated to be 1.0~1.8 × 10^−8^ per base pair (bp) per generation^2^. Germline mutation rates exhibit high heterogeneity across the genome, from single-nucleotide level to chromosome level^3^. Mutation rate is important for many genetic and evolutionary analyses, such as inferring population demographic histories^4^, detecting genomic regions undergoing natural selection^5^, and identifying disease-associated genetic variants^6^.

Despite its importance, constructing a fine-scale germline mutation rate map for a eukaryotic genome is particularly challenging. One main challenge is the rarity of DNMs in each generation, making it costly to obtain many high-quality DNMs using the gold standard family-based sequencing strategy (e.g., sequencing parents and offspring simultaneously). Many studies used within-species polymorphisms or interspecies divergence to estimate mutation rates, but a substantial fraction of variants at polymorphic or divergent sites are evolutionarily old and affected by natural selection and (or) nonadaptive processes such as GC-biased gene conversion^7^. Recent studies^8–10^ demonstrated that extremely rare variants derived from population polymorphism data can serve as a reasonable proxy for DNMs to predict mutation rates, ameliorating the condition of data insufficiency. Another challenge in estimating fine-scale mutation rates is the complex relationships between predictor variables and mutation rates. Adjacent nucleotides are significant predictors for SNV mutation rates of a focal nucleotide^3,11^, particularly the immediately 5’ and 3’ nucleotides. Nucleotides more distant from a focal site are also associated with mutation rate variation, though to a lesser extent^12,13^. Apart from sequence context, functional genomic features, such as DNA methylation, replication timing and recombination rate^14^, were reported to be associated with mutation rate variation and have been included in mutation rate modeling work^9,15–17^.

Existing models have several limitations. First, some models only considered a small number of adjacent nucleotides (typically not longer than 7-mer centered at the focal nucleotide), because modeling longer sequences requires more parameters that could make training difficult. Second, previous work mainly employed linear or generalized linear models to estimate mutation rates, but relationships between mutation rates and predictors could be nonlinear^18,19^ and more complicated. Third, some models required many mutations and (or) functional genomic features for training, which limits their use in species lacking published mutations and functional genomic data.

Deep learning methods have shown outstanding performance in solving difficult predictive problems^20^ and are increasingly used to address problems in genomics^21–26^. As the genomic sequence is the predominant factor for estimating mutation rates and many functional genomic features are correlated with the sequence, we reasoned that deep learning would be a promising approach to capturing various signals from genomic sequences to generate improved mutation rate maps.

With the above considerations, we developed a deep learning framework (named MuRaL, short for <u>Mu</u>tation <u>Ra</u>te <u>L</u>earner) to estimate single-nucleotide germline mutation rates across the genome with DNA sequences as input. Comprehensive assessment using human variant data showed that MuRaL-predicted mutation rates were highly correlated with observed mutation rates at different scales. Compared to current state-of-the-art models, MuRaL required much less training data and fewer sequenced individuals but exhibited improved performance. We further demonstrated that MuRaL can be easily applied to other species and the improved mutate rate estimates are beneficial for other analyses.

## Results

### Design of the MuRaL model

The MuRaL model has two main neural network modules (**Fig. 1; Supplementary Fig. 1;** see Methods), one for learning signals from local genomic regions (e.g., 5~10bp on each side of the focal nucleotide), the other for learning signals from expanded regions (e.g., 1Kb on each side). The main reason for having both modules is that local and distal sequences likely contribute to the mutability of a focal nucleotide in different ways, thus the signals in them might be better learned by different network architectures.

**Figure 1.**
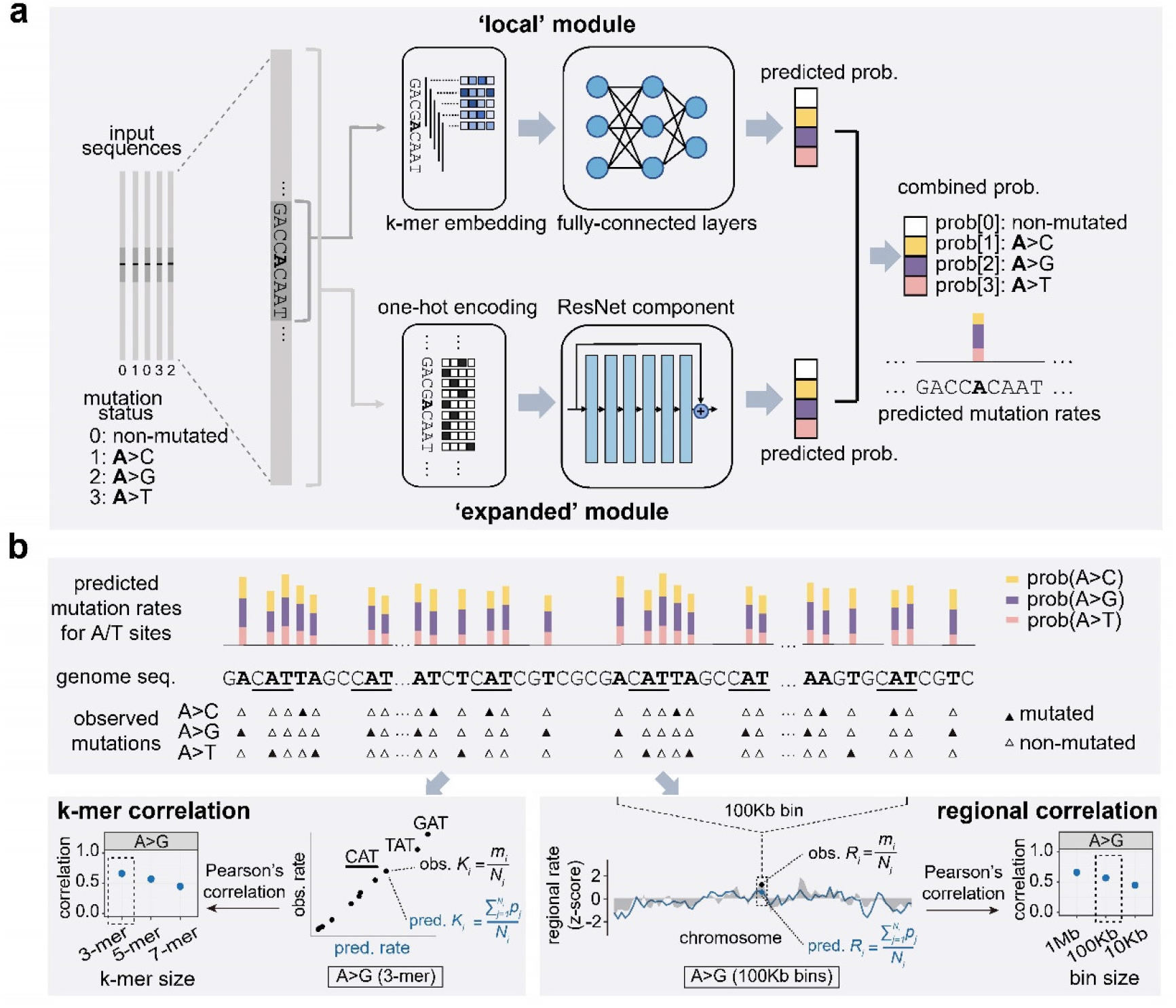
Schematics for the MuRaL model and evaluation strategies. (**a**) The model consists of a ‘local’ module and an ‘expanded’ module. In the ‘local’ module, the input sequence of the focal nucleotide (e.g., the bold ‘A’ in the figure) is split into overlapping k-mers which are then mapped into multi-dimensional vectors by the embedding layer. The multi-dimensional vectors are concatenated and passed to three fully-connected (FC) layers. The output of the ‘local’ module is a probability distribution generated by the softmax function over four predicted classes - non-mutated or one of three possible substitution mutations (e.g., A>C, A>G and A>T). In the ‘expanded’ module, the input sequence of an expanded region is one-hot encoded. The one-hot encoded matrix is considered as one-dimensional data with four channels and passed to a ResNet component. An additional FC layer and the softmax function following the ResNet component generate a probability distribution over four predicted classes, like that in the ‘local’ module. The probabilities of ‘local’ and ‘expanded’ modules are combined using equal weights (i.e., 0.5*P_local_ + 0.5*P_expanded_) to generate the combined probabilities. For training, the mutation status (bottom left corner) of each input sequence is also required. More details of the layers are provided in **Supplementary Fig. 1**. (**b**) An illustration of two key evaluation metrics - k-mer and regional correlations between observed and predicted averaged rates. The top panel shows the predicted mutation rates and observed mutations (rare variants or de novo mutations) at the nucleotide level. The bottom left panel shows how a 3-mer correlation (N[A>G]N) is calculated, and the highlighted 3-mer ‘CAT’ is also underlined in the top panel to indicate source positions. The bottom right panel shows how a regional correlation (A>G 100Kb bins) is calculated. More details of the two metrics are given in the Methods.

The ‘local’ module consists of a k-mer embedding layer and multiple fully-connected (FC) layers to learn signals from the input local sequence surrounding the focal nucleotide (**Fig. 1a; Supplementary Fig. 1**). The ‘expanded’ module has a series of convolutional neural network (CNN) layers, which form a typical Residual Network (ResNet) architecture, to learn signals from the expanded sequence space (**Fig. 1a; Supplementary Fig. 1**). The outputs of two modules are both a four-element vector of probabilities, which represent mutational probabilities of a focal nucleotide to another three possible nucleotides and the probability of being non-mutated. Probability vectors of two modules are combined using equal weights to form a vector of combined probabilities. The predicted single-nucleotide mutational probabilities were considered mutation rates for subsequent analyses (**Fig. 1a**).

Unlike many previous deep learning models in genomics, our model aimed to obtain reliable class probabilities rather than accurate classification (i.e., assign a sample to a specific class). As probabilities derived from neural networks are usually not well calibrated, we further applied a Dirichlet calibration method^27^ to obtain calibrated probabilities (**Supplementary Fig. 1**).

We trained separate models for A/T sites and C/G sites, respectively. Moreover, for genomes with exceptionally high mutation rates at CpG sites, we trained models for non-CpG C/G sites and CpG sites separately. The three were called, for short, AT, non-CpG and CpG models. We only considered mutational probabilities of substitutions in autosomes because other types of mutations and chromosomes have specific features that need to be modeled in a different manner. For analyses that needed to add up mutation rates of different site groups, we also scaled the mutation rates based on the mutation spectrum of DNMs (see Methods).

To evaluate the performance of different models, apart from cross-entropy losses in the validation data, we further considered two metrics - Pearson correlation coefficients between average observed and predicted mutation rates for k-mers and binned genomic regions, respectively (**Fig. 1b**, see Methods for more details). We considered k-mer and regional mutation rates for evaluation because observed mutations were very sparse across the genome, making it impossible to directly evaluate the accuracy of predicted mutation rates at single-nucleotide resolution.

### MuRaL can build effective models with relatively few variants from a moderate number of sequenced individuals

To evaluate the effectiveness and performance of MuRaL, we took advantage of the large-scale human variant data from gnomAD^15^ and generated multiple sets of rare variants for analysis (**Fig. 2a, Supplementary Table 1;** see Methods). We considered the reported issue^28^ that large sample sizes led to reduced proportions of observed CpG-related mutations (**Supplementary Fig. 2**). Unless specified elsewhere, we used the ‘1in2000’ data (allele frequency being 1/2000 after downsampling the total allele count to 2000) for training human AT and non-CpG models and the ‘5in1000’ data (allele frequency being ≤ 5/1000 after downsampling the total allele count to 1000) for training the CpG model (**Supplementary Table 2**). For detailed evaluation, we mainly used ‘10in20000’ rare variants (as observed mutations) for AT and non-CpG models, and ‘5in1000’ rare variants for CpG models because of their high variant densities (**Supplementary Table 1**).

**Figure 2.**
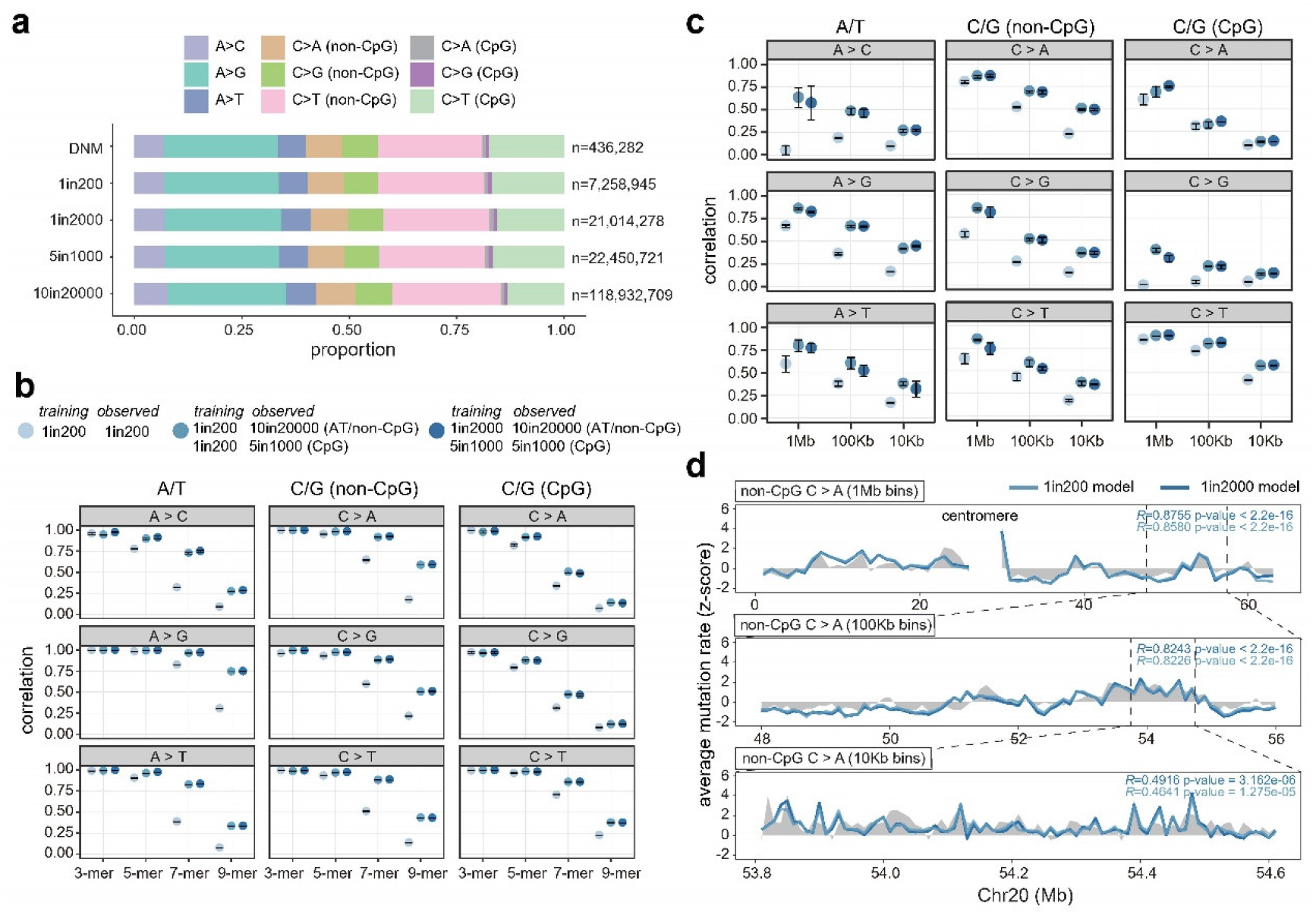
Comparison of MuRaL models trained with different rare variant data. (**a**) Mutation spectrums of multiple DNM or rare variant datasets, with numbers of total used variants on the right. (**b**) 3-, 5-, 7- and 9-mer correlations for different mutation types, based on predicted single-nucleotide mutation rates of Chr20 by different trained models and different observed mutation data (color scheme on top of the panel). Separate models were trained for A/T sites, non-CpG C/G sites and CpG sites, respectively. For each model, three best trials (out of ten) were used for calculating k-mer correlations, respectively. Dots and error bars indicate means and standard errors, respectively, for correlation coefficients derived from three trials. Decrease of correlations in longer k-mers is expected, because longer k-mers have fewer mutations per mutation subtype and thus average observed and predicted rates are more affected by sampling noise. (**c**) Regional correlations with bin sizes of 1Mb, 100Kb and 10Kb on Chr20 for different mutation types and different observed mutation data. The meanings of colors, dots and error bars are the same as that for panel **b**. (**d**) An example showing regional correlations for non-CpG C>A mutations at different scales on Chr20 for 1in200 and 1in2000 models, with grey shades indicating observed mutation rates (based on 10in20000 data) and colored lines for predicted rates. As predicted and observed regional mutation rates had different magnitudes, z-score normalization was applied for visualization. Mutation rates at centromeric regions were not available. Pearson correlation coefficients and p-values for shown regions are provided at the upper right corners. P-values of all correlation tests performed for panels **b** and **c** were provided in **Supplementary Data 1**.

Different network architectures and hyperparameters can affect model performance. By training experiments with different values, we chose the lengths of ‘local’ and ‘expanded’ regions to balance training efficiency and required resources (**Supplementary Fig. 3**). For most models in this work, we set the ‘local’ region length to be 15bp (7bp on each side on the focal nucleotide) and the ‘expanded’ region length to be 2001bp (1Kb on each side). Our ablation analyses revealed that two modules (‘local’ and ‘expanded’) of MuRaL have distinct advantages in learning mutability signals, indicating the necessity of both (**Supplementary Fig. 4**),.

Another important factor is the training data size. We found that increasing training data sizes continuously reduced the validation loss (**Supplementary Fig. 5**), but the computational burden increased substantially in turn. For the human genome, we used 500,000 mutated and 10,000,000 non-mutated sites for training each MuRaL model (**Supplementary Table 2**), unless specified otherwise.

As the number of mutations required for training MuRaL models was not large, we investigated MuRaL’s performance if training with rare variants from fewer sampled genomes. We tried training MuRaL models with ‘1in200’ rare variants (**Fig. 2a**; allele frequency being 1/200, i.e., occurring once in 100 diploid genomes). Because the ‘1in200’ data had a small number of rare variants and thus a low mutation density across the genome, we generally got low k-mer and regional correlations if using them for calculating observed mutation rates (**Fig. 2b, c**). However, with more dense rare variant datasets as observed mutations (‘10in20000’ and ‘5in1000’ data; **Fig. 2b, c**), the ‘1in200’ MuRaL models achieved much increased regional and k-mer mutation rate correlations, suggesting the high predictive performance of these models. This also implies that if only 100 human genomes are available, it is still possible to build reasonably good models by using the rare variants from the 100 individuals.

Moreover, we showed that AT and non-CpG models trained with singleton variants derived from the ‘1in200’ data had similar performance as those trained with the same amount of singleton variants from the ‘1in2000’ data (**Fig. 2b-d**). The mutation rate correlations of the CpG model trained with ‘1in200’ data were also close to that of the model trained with ‘5in1000’ data (**Fig. 2b, c**). These results further corroborated that MuRaL can train effective models with a relatively small number of rare variants from a moderate number of sequenced individuals.

### Other factors that might affect the performance of MuRaL

As read coverage can affect mutation calling and is usually accessible for mutation data, we further tried incorporating read coverage into MuRaL (**Supplementary Fig. 6;** see Methods). In well-covered regions, MuRaL models with coverage had similar performance as those without coverage (**Supplementary Fig. 7**). Since exonic regions usually experienced stronger selection, we also trained MuRaL models using only non-exonic rare variants, results of which also showed similar performance with respect to k-mer and regional correlations (**Supplementary Fig. 8;** see Methods). In addition, we tried training AT models using rare variants from only healthy controls in gnomAD or using rare variants after excluding singletons, the results of which showed little difference (**Supplementary Fig. 9;** see Methods).

We noticed that several chromosomes, such as Chr7, Chr9, Chr15 and Chr16, showed smaller regional correlations than others (**Supplementary Fig. 7**), likely due to their enrichment for recent segmental duplications^29^. The poor regional correlations of Chr8 was ascribable to under-estimated mutation rates in the region from 0Mb to 25Mb (**Supplementary Fig. 10**), a region reported to have a strikingly high mutation rate^30^. The relatively small learning space (2Kb) of MuRaL models may not efficiently capture distinct region-specific mutability signals in these complicated regions.

For some mutation types (e.g., CpG>GpG), because their numbers of training mutations were much smaller than others (**Supplementary Table 2**), related model parameters would be more susceptible to underfitting.

### MuRaL outperforms existing models

We compared MuRaL with several recently published models for estimating mutation rates across the human genome (**Fig. 4; Supplementary Table 3**). Among all models, MuRaL used the smallest number of training mutations (1.5 millions in total) and did not rely on any functional genomic data.

For genome-wide correlations between observed and predicted k-mer mutation rates, MuRaL, ‘Carlson 7-mer+features’ and ‘Carlson 7-mer’ models performed similarly and were much better than the other two models (**Fig. 3a**). Though for specific mutation types such as A>C and CpG>GpG, 5-mer and 7-mer mutation rate correlations of the Carlson models were slightly better than MuRaL, MuRaL showed better performance in almost all correlations for 9-mer mutation rates, probably because MuRaL considered sequence context beyond 7-mers. At the chromosome level, the patterns were similar to the genome-wide patterns (**Supplementary Figs 11-13**).

**Figure 3.**
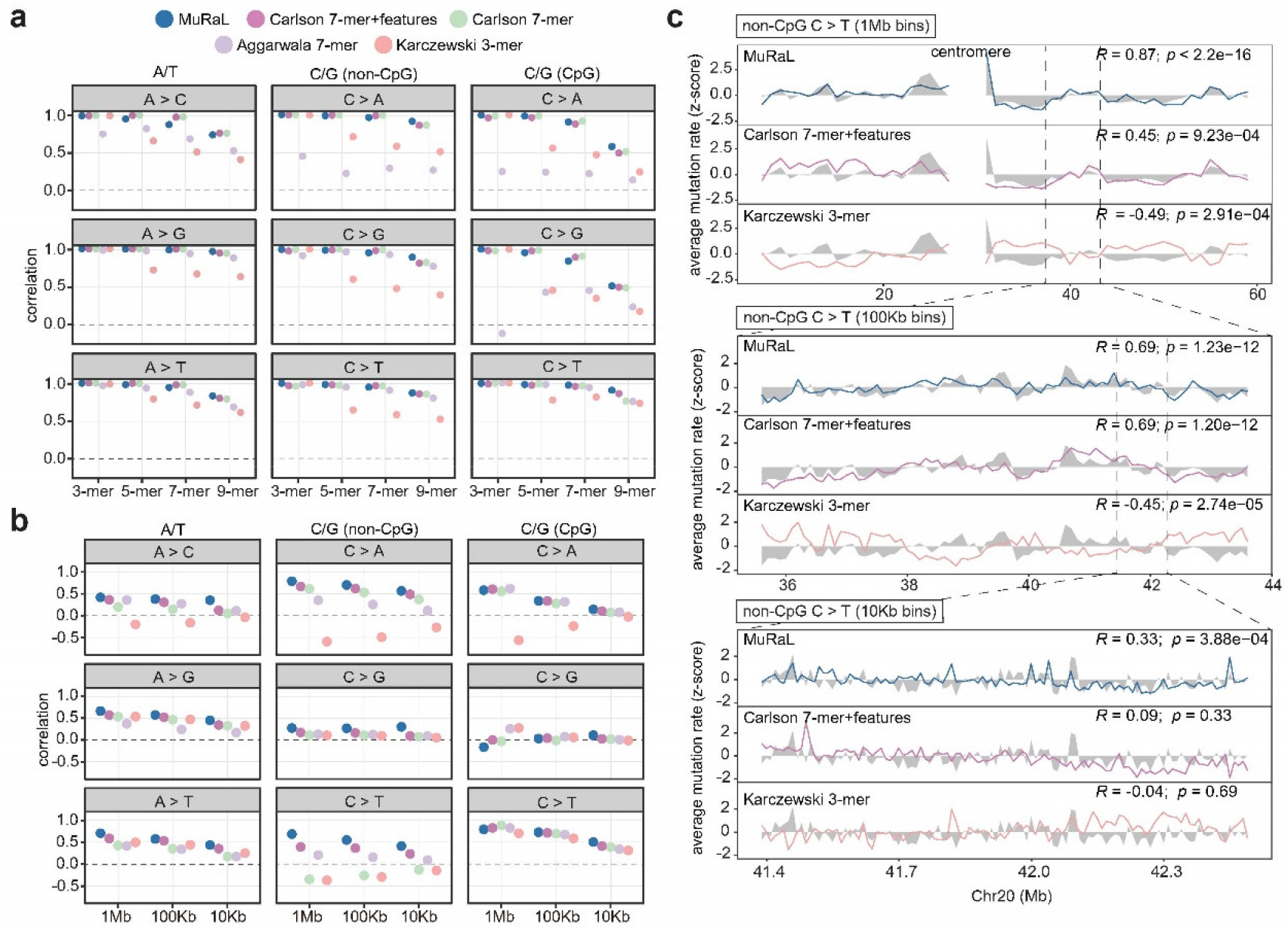
Comparison of MuRaL and existing models. (**a**) 3-, 5-, 7- and 9-mer mutation rate correlations for different mutation types, based on predicted single-nucleotide mutation rates of the autosomal genome by different models. The mutations for calculating observed mutation rates were ‘10in20000’ rare variants for AT and non-CpG models, and ‘5in1000’ rare variants for CpG models. (**b**) Regional mutation rate correlations with bin sizes of 1Mb, 100Kb and 10Kb on the autosomal genome for different mutation types. The color scheme was the same as that for panel **a**. (**c**) An example showing regional mutation rate correlations at different scales on Chr20 for three models (MuRaL, ‘Carlson 7-mer+features’ and ‘Karczewski 3-mer’), with grey shades indicating observed mutation rates and colored lines for predicted rates. As predicted and observed regional mutation rates had different magnitudes, z-score normalization was applied for visualization. Mutation rates at centromeric regions were not available. Pearson correlation coefficients and p-values for shown regions are provided at the upper right corners. P-values of all correlation tests performed for panels **a** and **b** were provided in **Supplementary Data 1**.

For correlations of regional mutation rates, MuRaL performed better than any other model at the genome-wide level (**Fig. 3b)**. In general, the superiority of MuRaL was more pronounced when the bin sizes were smaller, suggesting that MuRaL predicted improved mutation rates at finer scales compared to previous models. (**Fig. 3b, c; Supplementary Figs 11-13**). It is worth noting that, if aggregating three mutation types associated with the same reference base (e.g., merging A>C, A>G and A>T mutations), at the 1Kb scale MuRaL models still achieved regional correlations of ~0.3 (**Supplementary Fig. 14**). At the chromosome level, MuRaL performed best for most chromosomes and most mutation types (**Supplementary Figs 11-14**). For chromosomes that MuRaL had relatively low regional correlations, other models showed similar trends in most cases (**Supplementary Figs 11-14**). MuRaL also performed better than other models in exonic regions (**Supplementary Figs 15**).

Although previous models using only local sequence context (3-mers or 7-mers) generally had positive correlations for regional mutation rates, for specific mutation types (especially non-CpG C/G mutations), they had poor or even negative correlations (**Fig. 3b; Supplementary Figs 11-14**). This indicates that a short adjacent sequence cannot fully capture the signals related to the mutability of a focal nucleotide.

We noticed that coefficients of variation (CVs) of regional mutation rates from all the models were much smaller than that of observed regional mutation rates at different scales (1Mb, 100Kb and 10Kb; **Supplementary Fig. 16**). Although larger CVs of observed mutation rates could be partly due to sampling noise (especially for small bin sizes), the big differences between CVs of observed and predicted mutation rates suggested an aspect that needs to be improved in future.

### Training with DNMs and transfer learning

The number of published DNMs in humans is much smaller than that of rare variants. However, because MuRaL can work with relatively few training mutations, we tried training AT and nonCpG MuRaL models using 150,000 DNMs and the CpG model using 50,000 DNMs (**Supplementary Table 4**; see Methods).

Transfer learning is widely used in deep learning for scenarios in which the prediction tasks are similar but less training data is available. To study the effectiveness of transfer learning in the MuRaL framework, we trained transfer learning models with same DNMs, using pre-trained weights from aforementioned rare-variant models for parameter initialization. With independent validation DNMs (see Methods), we found that models with transfer learning achieved significantly lower validation losses than those without transfer learning (*ab initio* DNM models; **Fig. 4a**). Transfer learning models also showed generally better k-mer and regional correlations when using DNMs as observed mutations (**Fig. 4b, c**).

**Figure 4.**
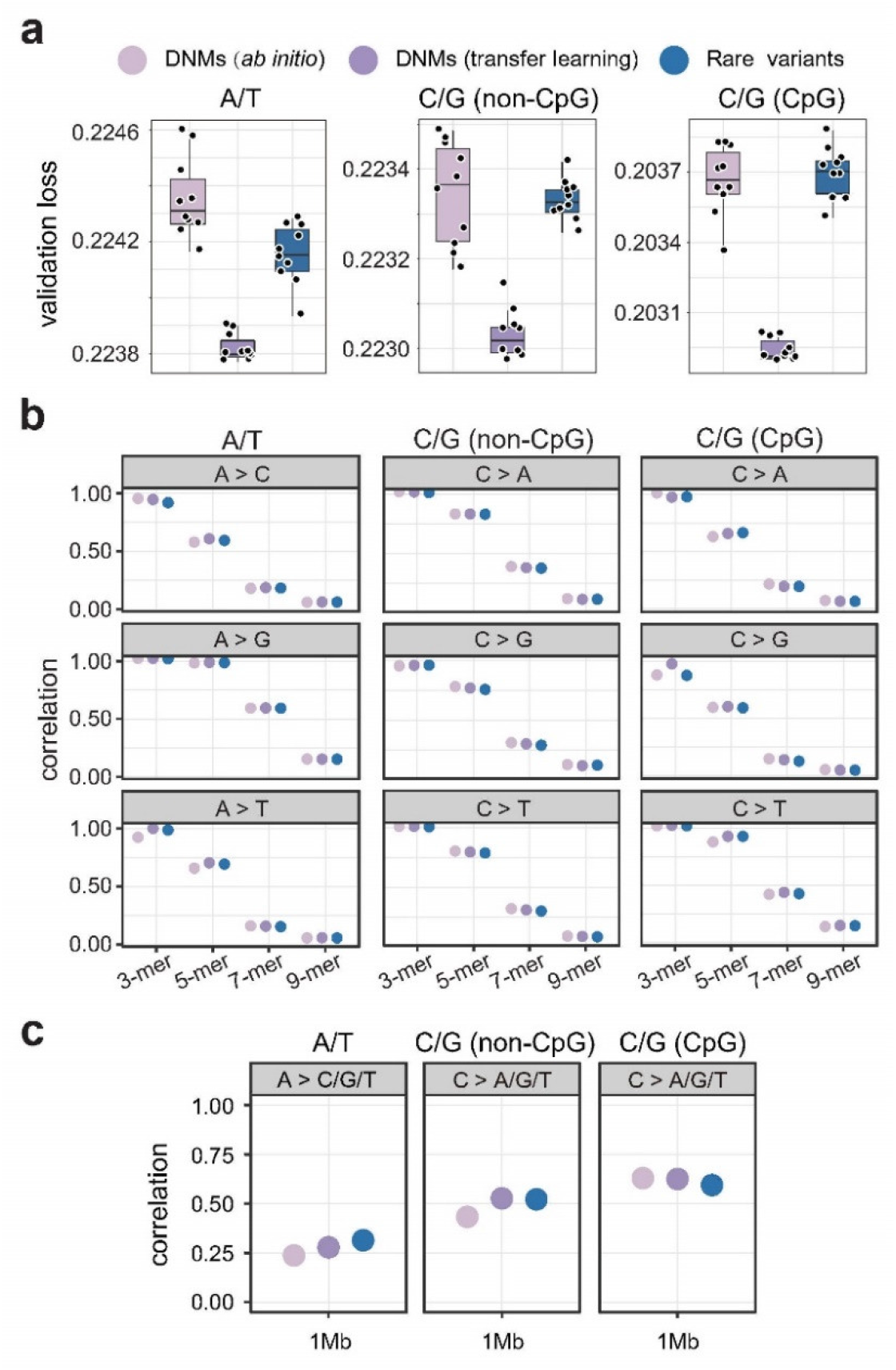
Training DNM models and transfer learning. (**a**) Validation losses on the validation DNMs for three types of models: DNM *ab initio* models, DNM transfer learning models, and rare-variant models. For each model, the lowest loss (mean cross-entropy loss) for each of ten trials was used to generate the boxplots. (**b**) 3-, 5-, and 7-mer mutation rate correlations for different mutation types, based on predicted single-nucleotide mutation rates on human Chr1. Colors depict three types of models like that in panel **a**. For each model in panel **a**, the best trial with the lowest validation loss was used for predicting mutation rates on Chr1. The mutations for calculating observed mutation rates were human DNMs. (**c**) Regional mutation rate correlations with a 1Mb bin size on Chr1. The predicted mutation rates of multiple mutation types (e.g. A>C/A>G/A>T) were aggregated for calculating regional correlations, as some mutation types had very few observed DNMs in the data. Smaller bin sizes were not assessed due to few DNMs. Colors depict three types of models like that in panel **a**. P-values of all correlation tests performed for panels **b** and **c** were provided in **Supplementary Data 1**.

Furthermore, we computed validation losses of the validation DNMs using the rare-variant models described in previous sections. Compared to the *ab initio* DNM models, the rare-variant models achieved lower or similar validation losses for all three categories of mutations **(Fig. 4a**). When looking at k-mer and regional correlations, rare-variant models generally performed better than DNM *ab initio* models, and similarly to the DNM transfer learning models (**Fig. 4b, c**). This indicates that if DNMs are unavailable, we can reasonably use mutation rates predicted by rare-variant models to approximate *de novo* mutation rates.

When DNMs are available but limited, it might be beneficial to train transfer learning models with DNMs using pre-trained weights of rare-variant models. However, we note that DNMs collected from different studies could introduce biases, which need to be considered for transfer learning. For example, we found that our collected DNMs were substantially depleted in low-complexity regions and segmental duplications relative to rare variants (**Supplementary Fig. 17**), probably due to conservative variant calling procedures.

### Generating mutation rate maps for other species

We further applied MuRaL to estimate mutation rates for three other species. For species that are evolutionarily close to humans, their genomes have high sequence similarities with the human genome, and many mutational processes are likely shared between them. Hence transfer learning can be leveraged for those species. The rhesus macaque (*Macaca mulatta)* is a close relative of humans and a widely used model organism. We trained *ab initio* MuRaL models as well as transfer learning models for *M. mulatta* using the rare variants from a dataset of 853 individuals^31^. The training data size of transfer learning models was 30% of that for *ab initio* models (**Supplementary Table 5;** see Methods). Transfer learning models showed similar performance to that of *ab initio* models (**Fig. 5a, b**), though transfer learning models used less training data and computation time (**Supplementary Fig. 18**).

**Figure 5.**
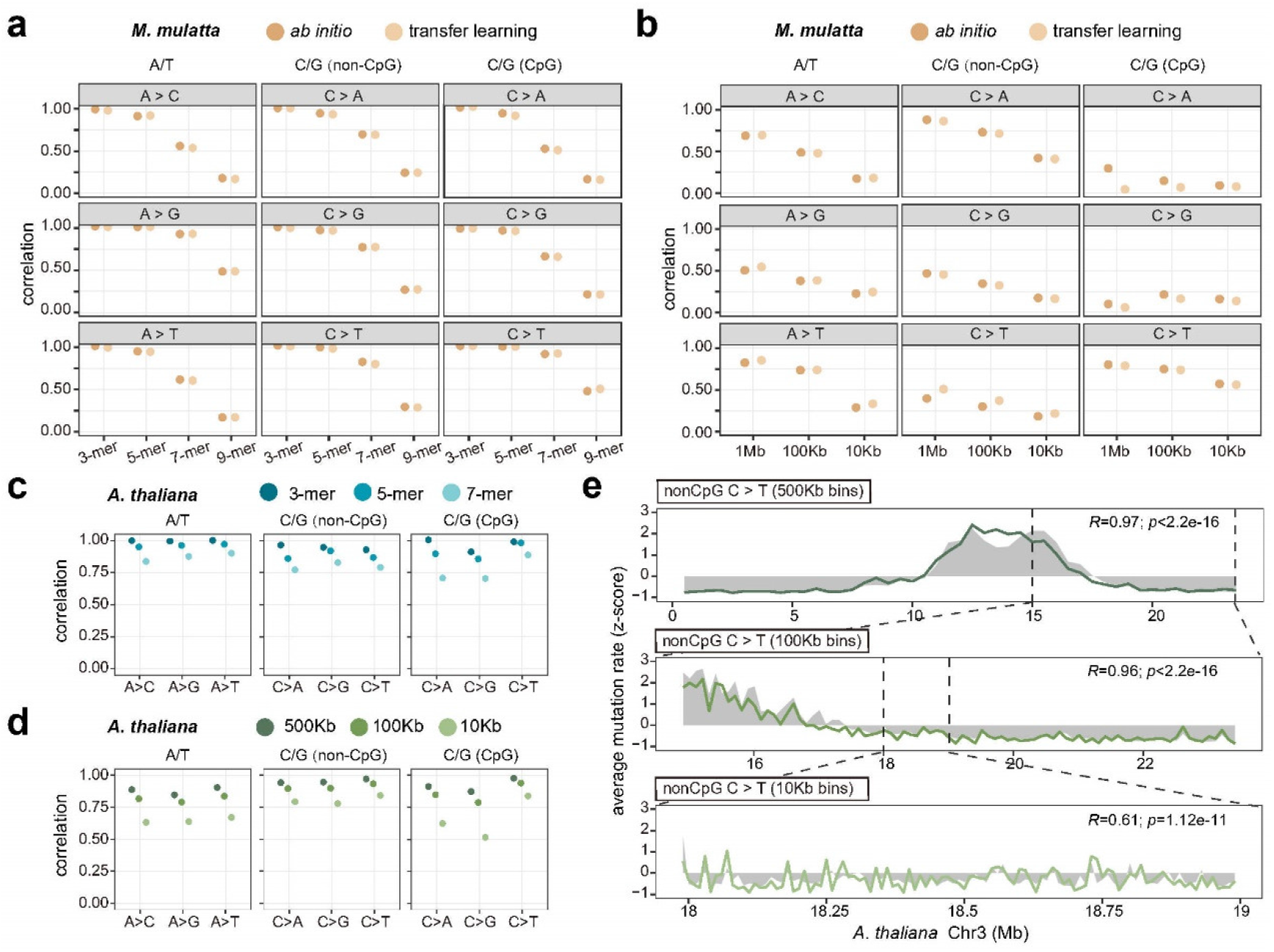
Application of MuRaL to other species. (**a**) 3-, 5-, 7- and 9-mer mutation rate correlations for different mutation types, based on predicted single-nucleotide mutation rates on rheMac10 Chr20 by two kinds of models for *M. mulatta: ab initio* models and transfer learning models. Rare variants of *M. mulatta* were used for calculating observed mutation rates. Separate models were trained for A/T sites, non-CpG C/G sites and CpG sites, respectively. (**b**) Regional mutation rate correlations with bin sizes of 1Mb, 100Kb and 10Kb on rheMac10 Chr20 for different mutation types. (**c**) 3-, 5-, and 7-mer mutation rate correlations for different mutation types, based on predicted single-nucleotide mutation rates for the *A. thaliana* genome. Rare variants of *A. thaliana* were used for calculating observed mutation rates. Separate models were trained for A/T sites, non-CpG C/G sites and CpG sites, respectively. (**d**) Regional mutation rate correlations with bin sizes of 500Kb, 100Kb and 10Kb for the *A. thaliana* genome. (**e**) An example showing regional mutation rate correlations at different scales on Chr3 for *A. thaliana*, with grey shades indicating observed mutation rates and colored lines for predicted rates. As predicted and observed regional mutation rates had different magnitudes, we applied z-score normalization for visualization. Pearson correlation coefficients and p-values for shown regions are provided at the upper right corners. P-values of all correlation tests performed for panels **a-d** were provided in **Supplementary Data 1.**

We also trained *ab initio* MuRaL models for two model organisms that are evolutionarily distant to humans - *Drosophila melanogaster* and *Arabidopsis thaliana*. As these genomes (<200Mb) are much smaller than the human genome, we used only 100,000 mutations for training each of the models (**Supplementary Tables 6-7**; see Methods). Despite a relatively small amount of training data, predicted results of trained models suggested that MuRaL worked well in these species in terms of k-mer and regional correlations (**Fig. 5c-e; Supplementary Fig. 19**). For example, for *A. thaliana*, at the scale of 10Kb bins, regional correlations were >0.5 for all mutation types (**Fig. 5d, e**), indicating high effectiveness of our method in this species. For *D. melanogaster*, we used singleton variants from only 205 inbred lines for training and still obtained promising results (**Supplementary Fig. 19**), further demonstrating that MuRaL can be applied to scenarios with a relatively small number of sequenced genomes.

### Using MuRaL-predicted mutation rates for other analyses

Based on MuRaL-predicted mutation rates, we generated meta-gene mutation rate plots for regions around coding genes of the human genome. We found that promoter regions exhibited notably elevated average mutation rates (**Fig. 6a**), as reported previously^32^, indicating that they are mutational hotspots. The high-resolution MuRaL mutation rates allowed us to cluster genes based on mutation rate profiles and plot fine-scale heatmaps (**Fig. 6b,c**), which could not be done with few observed variants. Interestingly, K-means clustering with MuRaL mutation rates stratified the genes into three distinct clusters which were enriched for different Gene Ontology (GO) categories (**Fig. 6d; Supplementary Fig. 20**), and such stratification was reflected by meta-gene plots of observed rare variants (**Fig. 6b**). Notably, the gene cluster with elevated mutation rates in both gene bodies and flanking regions was enriched with various development-related GOs (**Fig. 6d**), suggesting that many developmental genes are subject to high mutational burden. This might have important implications in evolution and disease. It is worth noting that clustering using ‘Carlson 7-mer+features’ and ‘Karczewski 3-mer’ rates did not generate stratifications compatible with observed variants (**Supplementary Fig. 21**).

**Figure 6.**
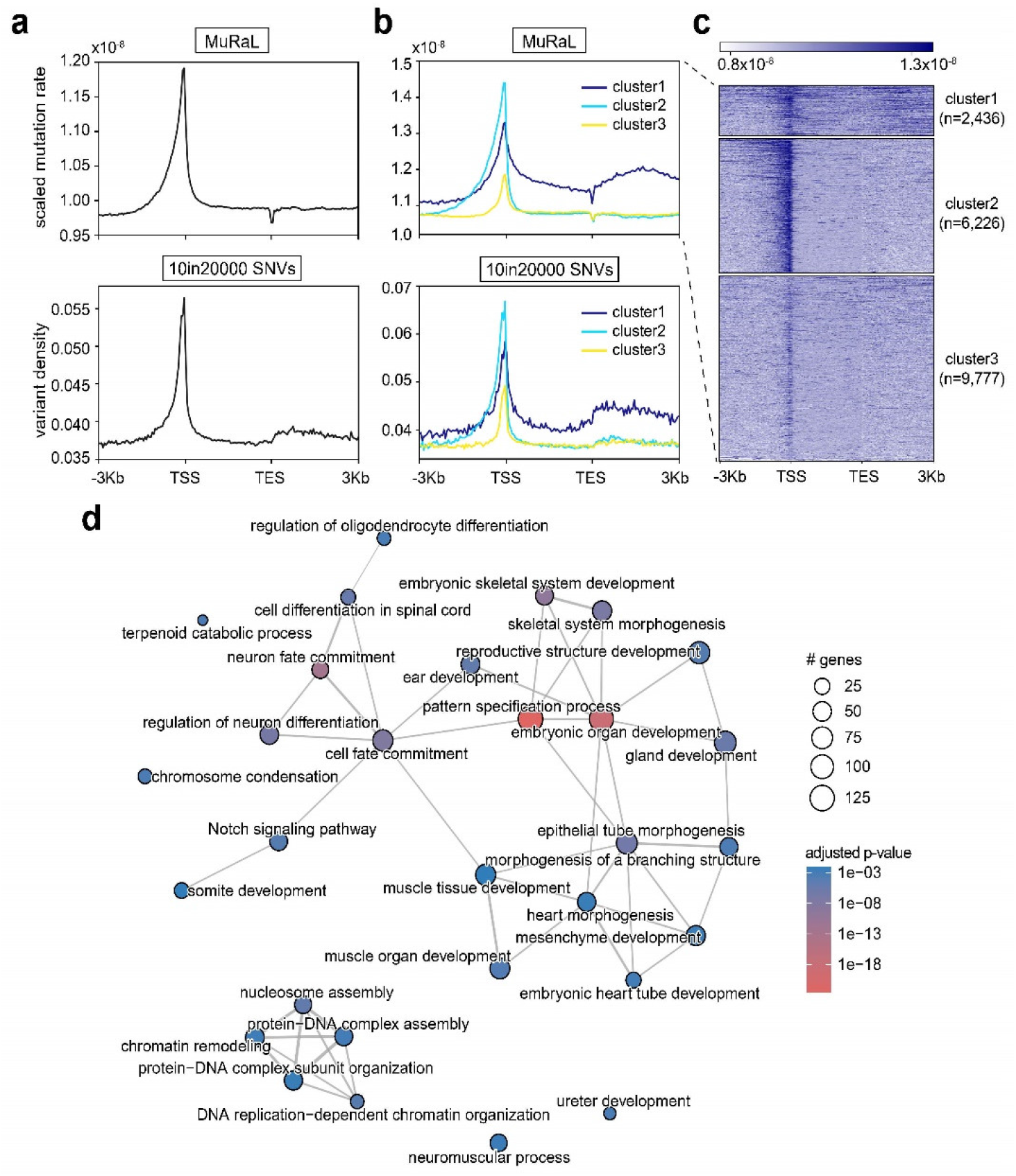
Clustering human coding genes based on mutation rate profiles. (**a**) Meta-gene mutation rate plots with a bin size of 50bp for regions around human coding genes. Gene bodies were scaled to 3Kb. The upper was based on scaled MuRaL mutation rates (see Methods), and the lower based on ‘10in20000’ rare variants. TSS, transcription start site; TES, transcription end site. (**b**) Meta-gene plots of three gene clusters after performing K-means clustering (K=3) with MuRaL mutation rates (same data in panel **a**). The same clusters were used for generating meta-gene plots for ‘10in20000’ data (lower panel). (**c**) The heatmap of MuRaL mutation rates of three clusters in panel **b**, with the number of genes in clusters given in brackets. (**d**) Enriched GO terms of the ‘cluster1’ genes of panel **c**.

As another example, based on MuRaL mutation rates and gnomAD variants, we generated mutational tolerance scores for sliding 500bp windows across the human genome using the depletion rank (DR) method in a recent UK Biobank study^33^ (see Methods). We found that regions with lowest DR scores were enriched for pathogenic variants in ClinVar, and the enrichment was significantly higher when using MuRaL mutation rates for DR calculation than using Carlson mutation rates (Fisher’s exact test, *p* < 2.2e-16 for lowest 10% DR, **Supplementary Fig. 22**). This indicates that DR scores derived from MuRaL mutation rates are informative for prioritizing disease-related variants.

## Discussion

Estimation of sequence mutation rates in the genome can be traced back to the very early period of molecular evolution research^34^. Being hindered by the lack of genomic data, early work only obtained rough estimates of mutation rates for specific genes or genomes. In past few years, by taking advantage of large-scale variant data, several methods have been proposed to infer fine-scale mutation rates, but there was much room for improvement.

In this work, we developed the MuRaL framework to address the challenge. Compared with the previously best-performed model by Carlson et al.^9^, MuRaL learned signals from a much larger sequence space, required fewer training mutations and did not rely on functional genomic data. Importantly, the relaxed requirements for training data open opportunities for generating fine-scale mutation rate maps for other species. Our successful application of MuRaL to four representative species for primates, insects and plants demonstrated its high applicability. For non-human species (especially non-model species), we note that available reference genomes and variants could have more artifacts, including those related to genomic complexity, sequencing technologies, variant calling pipelines and population histories. One should be more cautious when performing data preprocessing and model training for non-model species.

The generated mutation rates and software are relevant for many analyses. For example, we demonstrated that the high-resolution mutation rates are useful for identifying genes or regions with high mutability in the human genome. The predicted mutation rates can help improve previous methods of calculating mutation intolerance scores^11,15,33,35^ for prioritizing disease candidate genes or variants, as exemplified by the re-calculated DR scores in our work. The predicted mutation rates should also be informative for detecting regions undergoing selection or introgression in recent evolution. As there are already many sequenced genomes for different human populations, it should not be difficult to generate population-specific mutation rate maps with MuRaL. Comparison of mutation rate maps between populations or species can advance our understanding of mutation rate evolution as well as underlying mutational mechanisms.

Several aspects can be investigated or improved in the near future. Some genomic regions, such as those overlapping recent segmental duplications on human chromosome 8, showed relatively poor mutation rate estimates. To address this, using longer input sequences and (or) function genomic data for training may offer more signals, at the expense of more computational load. Given the rapid development of deep learning hardware, we believe computational load will become a minor obstacle soon. The MuRaL framework can be extended to estimate mutation rates for sex chromosomes and organelle genomes, though more specific assessments are required. Similar models could also be developed to predict fine-scale mutation rates for other mutations such as small insertions and deletions.

To our knowledge, this is the first time that deep learning is used to estimate fine-scale germline mutation rates. Unlike many deep learning models in genomics designed for typical classification or regression problems, our method aimed to predict accurate class probabilities. This work provided an exemplary case for addressing similar problems in genomics. How to obtain reliable class probabilities in deep learning models is still a hot research topic in computer science^36^. The smaller CVs of predicted regional mutation rates than that of observed rates might be partly due to the large class imbalance in the training data, which often causes issues in deep learning models. Future progress on these topics in computer science may help improve our model.

In summary, we developed a generalizable framework for inferring fine-scale mutation rates of genomes, which can facilitate and stimulate future research in related fields.

## Methods

### Design of the MuRaL model

Previous studies revealed that adjacent nucleotides of a specific site predominantly affect its mutation rate and properties of a larger sequence context (e.g., GC content, replication timing) are also associated with mutation rate variation. As local and distal sequences likely affect mutation rates in different ways, we constructed two different neural network modules to learn the signals from the two aspects. One module (termed ‘local’ module) was designed for learning signals from a local sequence of the focal nucleotide, the other (termed ‘expanded’ module) for learning signals from an expanded sequence (**Supplementary Fig. 1**).

The ‘local’ module consists of an embedding layer and three fully-connected (FC) layers to learn signals from the sequence. We used k-mer embedding because it was reported to offer benefits for deep learning models in genomics^37^. The input local sequence was firstly split into overlapping k-mers and the embedding layer maps these k-mers into multi-dimensional vectors. The multi-dimensional vectors from an input sequence were then concatenated to form the input for subsequent two hidden layers and one output layers. For each FC layer, ReLU (Rectified Linear Unit) activation function was used with the output of the FC layer, followed by batch normalization and dropout layers which would facilitate learning and avoid overfitting. The outputs of the ‘local’ module were probabilities of four-class classification of input samples, representing the probabilities of the focal nucleotide mutated to another three possible nucleotides or being non-mutated.

In the ‘expanded’ module, the sequence of an expanded region was first converted into four-dimensional vectors using one-hot encoding. Regarding one-hot encoding, each of the four bases (‘A’, ‘C’, ‘G’ and ‘T’) was converted to a four-element vector, in which all the elements were 0 except for one (e.g., ‘A’ converted to the vector [1, 0, 0, 0], ‘C’ converted to [0, 1, 0, 0]). The resulting matrix of the sequence was considered as one-dimensional data with four channels and then passed to a series of one-dimensional convolutional neural network (CNN) layers. The CNN layers were designed following a typical Residual Network (ResNet) architecture^38^. Two ResNets were employed for learning medium-scale and large-scale sequence signals, respectively. Each of the two ResNets was followed by a FC layer to produce probabilities of four-class classification of input samples, which were then combined with equal weights to form the probabilities of the ‘expanded’ module.

Finally, probabilities of ‘local’ and ‘expanded’ modules were combined using equal weights to form a vector of combined probabilities.

The key hyperparameters in the MuRaL model are the length of local sequences, the length of expanded sequences, the length of k-mers in the embedding layer, sizes of two hidden FC layers in the ‘local’ module, the kernel size and the number of channels for convolutional networks in the ‘expanded’ module (see **Supplementary Fig. 1**).

### Model implementation

We implemented the MuRaL model with PyTorch framework^39^, along with APIs from pybedtools^40^ and Janggu^41^. For model training, we used the cross-entropy loss function and the Adam optimizer^42^ for learning model parameters, and employed Ray Tune^43^ to facilitate hyperparameter tuning (**Supplementary Fig. 1**). The scheduler ‘ASHAScheduler’ in Ray Tune was used to coordinate trials and execute early stopping before reaching the specified maximum number of training epochs (e.g., 10), which can substantially reduce the training time. The mean cross-entropy loss of the validation sites (i.e., validation loss) was calculated at the end of each training epoch. We further set a stopping rule to terminate a trial if three consecutive epochs did not obtain a validation loss smaller than the current minimum validation loss. The ‘learning rate’ and ‘weight decay’ of Adam optimizer were two hyperparameters that could affect the learning performance significantly. We used Ray Tune to run trials with different values for the two hyperparameters. To have better convergence, we used the learning rate scheduler ‘lr_scheduler.StepLR’ in PyTorch to decay the learning rate after a number of steps by a specified factor.

### Human mutation data for model training and evaluation

#### Rare variants from gnomAD

Rare variants generally arose recently in the genome and were less affected by natural selection and nonadaptive evolutionary processes than common variants. Previous studies^8–10^ have established that rare variants can be used for estimating mutation rates. For model training and evaluation, we took advantage of the gnomAD database (v2.1.1) which contained genetic variation of 15,708 whole genomes^15^. Only single nucleotide substitutions in autosomes were considered, as other mutation types and sex chromosomes have specific features that need to be modeled separately. We extracted rare variants from gnomAD to approximate DNMs.

When the sample size is as large as that of gnomAD, some mutation types (e.g., CpG>TpG) with high mutation rates could be close to saturation, and the probability of multiple independent mutations (recurrence) at a same position increases. Therefore, we downsampled the gnomAD data into a specified total allele count using a hypergeometric distribution (see the probability density function below), and generated the random alternative allele counts from the hypergeometric distribution:

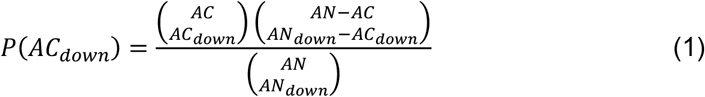

where AN and AC are the total allele count and the alternative allele count in original data, respectively, AN_down_ and AC_down_ are the total allele count and the alternative allele count after downsampling, respectively. For each polymorphic position, given values of AN, AC and AN_down_, we generated a random number for AC_down_ using the hypergeometric distribution of equation (1). We then extracted the variants with a specific alternative allele frequency (i.e., AC_down_/ AN_down_) in the downsampled data to form the rare variant datasets for subsequent analyses.

First, we downsampled the gnomAD data to total allele counts of 200, 2000 and 7000 (corresponding to 100, 1000 and 3500 diploid genomes) respectively and extracted the singleton variants (corresponding to AFs of 1/200, 1/2000 and 1/7000) in the three downsampled datasets. The three rare variant datasets were named ‘1in200’, ‘1in2000’ and ‘1in7000’, respectively. We considered the sample size of 7000 because Carlson et al.^9^ recently used singleton variants from a population of ~3500 individuals for modeling mutation rates. In addition, Karczewski et al.^15^ used variants with ≤5 copies in a downsampled set of 1000 haploid genomes for mutation rate estimation, so we also did downsampling for the sample size of 1000 and extracted the variants of AC_down_ ≤5 (termed ‘5in1000’). To increase the mutation density for smaller-scale evaluation, we further did downsampling for the sample size of 20000 and extracted the variants of AC_down_ ≤10 (termed ‘10in20000’). For SNVs with two or more different alternative alleles in the original gnomAD data, we did hypergeometric sampling for each alternative allele. In the downsampled dataset, if more than one alternative allele satisfied the rare variant criterion for a specific position, only one alternative allele was randomly selected for downstream analyses (i.e., not allowing multiple rare variants at one position). The numbers of rare variants in different datasets were summarized in **Supplementary Table 1**.

#### De novo mutations

We collected DNMs from the gene4denovo database^44^ for analysis. Because some data sources in gene4denovo database contributed only a small number of DNMs and different studies used distinct methods for variant calling, we used only the DNMs from three large-scale studies^45–47^ for our analysis, which consisted of 445,467 unique *de novo* SNVs.

To check whether the extracted rare variants can well represent properties of DNMs, we compared mutation spectra of rare variants and DNMs. We counted the occurrences of 1-mer and 3-mer mutation types for each dataset and calculated the relative proportion of each mutation type in the specific dataset. We found that mutation spectra of rare variants were highly similar with that of DNMs (**Supplementary Fig. 2**). However, when the sample size increased, the proportion difference in CpG>TpG mutation subtypes between rare variants and DNMs became larger (**Supplementary Fig. 2**). This was not surprising as mutation rates of CpG>TpG mutation subtypes were highest among all. Because genomic regions with too low or too high read coverage could have a high probability of false positives/negatives for mutation calls, we utilized the coverage information from gnomAD to exclude the positions with too low or too high read coverage. The genome-wide mean coverage per individual in the gnomAD data was 30.5, and genomic positions within the coverage range of from 15 to 45 (2,626,258,019 bp in autosome retained and considered as high-mappability sites) were used for downstream analyses.

#### Training and validation data for human MuRaL models

For the human data, we trained separate models for A/T sites, non-CpG C/G sites and CpG C/G sites, respectively. For training each MuRaL model, we randomly chose 500,000 mutations and 10,000,000 non-mutated sites. During training, we used an independent validation dataset consisting of 50,000 mutations and 1,000,000 non-mutated sites for evaluating training performance. The configuration of key hyperparameters for human MuRaL models was provided in **Supplementary Table 8**. As shown above, rare variants derived from a large sample of population could lead to depletion of mutation types of high mutability. On the other hand, rare variants derived from a small sample were relatively ancient and more affected by selection or other confounding processes. To balance the two constraints, we used the ‘1in2000’ data for training models of A/T sites and non-CpG C/G sites in human. Because ‘1in2000’ data showed more depletion of CpG>TpG mutations than ‘1in200’ and ‘5in1000’ data, it was not the ideal data for training the model of CpG sites. Although both ‘1in200’ and ‘5in1000’ datasets were rare variants of AC_down_/AN_down_≤0.005, we chose ‘5in1000’ data for training the CpG model because of its larger number of mutations. For detailed model evaluation, we mainly used ‘10in20000’ rare variants for A/T and non-CpG models, and ‘5in1000’ rare variants for CpG models due to their high mutation densities (**Supplementary Table 1**).

### Calibrating and scaling predicted probabilities

The main aim of our work is to obtain reliable class probabilities rather than accurate classification (e.g., predicting ones or zeros). Because probabilities from neural networks are usually not well calibrated, after training a MuRaL model, we applied a Dirichlet calibration method^27^ on the output combined probabilities to obtain better calibrated probabilities. Parameters of a Dirichlet calibrator were estimated by fitting the calibrator to the predicted probabilities of the validation data. Metrics such as Expected Calibration Error (ECE), classwise-ECE and Brier score^27^ were used for evaluating the performance of Dirichlet calibration. By comparing predicted mutation rates of validation data before and after calibration, we found that Dirichlet calibration indeed resulted in better ECE, classwise-ECE and Brier scores (**Supplementary Fig. 23**), although the improvements appeared to be relatively small. Small values of ECE and classwise-ECE scores before calibration (**Supplementary Fig. 23**) suggested that the original predicted mutation probabilities were already quite well calibrated in terms of such metrics.

The absolute values of above combined probabilities were not mutation rates per bp per generation. To obtain a mutation rate per bp per generation for each nucleotide, we may scale the calibrated probabilities using previously reported genome-wide DNM mutation rate and spectrum per generation. The scaling factor (*f^mut^* in the equations below) for a specific site group (e.g. A/T sites) can be calculated using following equations:

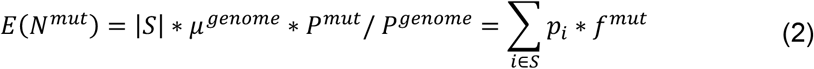

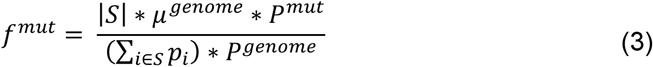

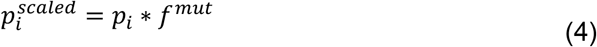

where *E*(*N^mut^*) is the expected number of mutations of a specific site group, *μ^genome^* is the per-base per-generation mutation rate obtained from DNM data, |*S*| is the number of considered genomic sites (*S*) for calculating scaling factors, *P^mut^* is the proportion of the specific mutation types in the DNM dataset, and *P^genome^* is the proportion of the specific site group in the genome. *p_i_*, and 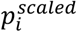 are single-nucleotide mutation probabilities before and after scaling. For the human genome, we used the *μ^genome^* of 1.2 x 10^−8^ (according to a recent study^48^) and validation sites of MuRaL models (*S*) to calculate the scaling factors, and then generated scaled mutation rates. We note that whether to do or not do this scaling does not affect the calculation of k-mer and regional mutation rate correlations described below.

### Extending MuRaL with read coverage

Read coverage (or read depth) of alignments can affect variant calling and thus observed mutation densities. We tried extending the MuRaL model to incorporate the coverage information (**Supplementary Fig. 6**). We used the pre-compiled coverage track from gnomAD (v2.1.1). In the ‘local’ module, we calculated mean coverage of the local sequence of the focal nucleotide, and added it as an additional element to the concatenated vector of embeddings of the local sequence. In the ‘expanded’ module, we extracted a coverage vector for the nucleotides of the expanded sequence, and merged it with the one-hot encoded matrix of the expanded sequence to form a five-channel input for subsequent convolutional networks. Such a design can also easily incorporate other genome-wide tracks (e.g., replication timing, recombination rate, etc.) to extend the MuRaL model.

### Correlation analysis of k-mer mutation rates

We classified mutations into six mutation types according to the reference and alternative allele: A>C, A>G, A>T, C>A, C>G, and C>T. Mutations with reference nucleotides T and G were reverse-complemented to that with A and C, respectively. For each mutation type, the k-mer subtypes were defined by the upstream and downstream bases flanking the variant site. For example, there are four possible bases at both the upstream −1 position and downstream +1 position, respectively, so there are 6 × 4^2^ = 96 3-mer subtypes, 16 3-mer subtypes for each basic mutation type. Similarly, for 5-mers and 7-mers, there are 6 × 4^4^ = 1,536 and 6 × 4^6^ = 24,576 subtypes respectively. In some analyses, we also considered 9-mers (6 × 4^8^ = 393,216 9-mer subtypes) if the number of mutations was large enough. For example, for the mutation type A>G, G[A>G]C and AG[A>G]CT are a 3-mer subtype and 5-mer mutation subtype associated with it respectively.

For the *i*th k-mer subtype, we calculated the observed mutated rate 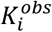 and the predicted mutation rate 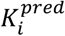 in the considered regions:

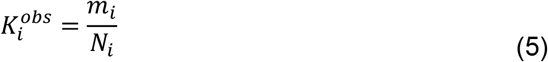

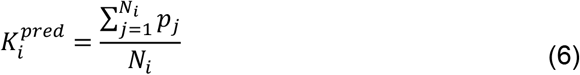

where *m_i_* is the observed number of mutated sites belonging to that k-mer subtype, *N_i_* is the total number of sites harboring the reference k-mer motif (e.g., all AAT 3-mers for the subtype A[A>C]T), and *p_j_* is the predicted mutation probability of the *j*th valid site.

Based on the calculated observed and predicted k-mer mutation rates, we can calculate the Pearson correlation coefficient for any set of k-mer subtypes (one subtype as a datapoint). For example, we can calculate a correlation coefficient for 16 3-mer subtypes associated with the mutation type A>C in a specific chromosome. Note that for CpG sites, there are only four 3-mer subtypes for a basic mutation type as the +1 position is fixed to be ‘G’. The Dirichlet calibrated mutation rates were used for calculating k-mer mutation rates unless otherwise specified.

### Correlation analysis of regional mutation rates

Since it was impossible to evaluate accuracy of predicted mutation rates on the single-nucleotide level, we compared average mutation rates in binned regions and calculated correlations between observed and predicted regional mutation rates to evaluate the performance of predicting models.

More specifically, for a specific mutation type, we calculated the observed and predicted mutation rates as below.

First, we divided a specified region (e.g., a chromosome) into non-overlapping bins with a given bin size (e.g. 10kb, 100kb, etc.). For the *i*th binned region, we calculated the observed mutated rate 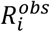 and the predicted mutation rate 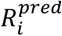,

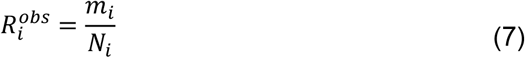

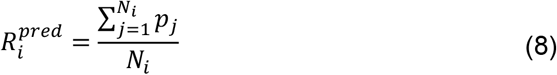

where *m_i_* is the number of observed mutations of the specific mutation type (e.g. A>C), *N_i_* is the total number of sites with same base as the reference base of that mutation type (e.g. all A/T sites for the mutation type A>C), and *p_j_* is the predicted mutation probability of the *j*th valid site in the binned region.

Based on the calculated observed and predicted regional mutation rates, we can calculate the Pearson correlation coefficient for any set of binned regions (one bin as a datapoint). For example, we can calculate the correlation coefficient for regional mutation rates of the mutation type A>C in all 100kb bins in a chromosome. As there are gaps and low-mappability regions in the genome, to avoid using regions with few valid sites for correlation analysis, we only used the bins that fit the criterion *N* > 20% * *N_median_*, where *N_median_* is the median of numbers of valid sites in all bins for a chromosome. The Dirichlet calibrated mutation rates were used for calculating regional mutation rates unless otherwise specified.

### Comparison of models with different network architectures

To see how the ‘local’ and ‘expanded’ modules contribute to the model, we considered three models – the ‘local-only’ model containing only the ‘local’ module, the ‘expanded-only’ model containing only the ‘expanded’ module and the full model with both modules. We trained the models with a training dataset of 500,000 mutated and 10,000,000 non-mutated sites randomly selected from autosomes. For each network architecture, ten trials were trained with Ray Tune. A validation dataset consisting of 50,000 mutated and 1,000,000 non-mutated sites was used to compare performance of three architectures based on the validation losses of trained trials. We also used the best trained trial with lowest validation loss for each of three architectures to predict mutation probabilities on the whole chromosome of human Chr20, results of which were then passed to compute k-mer/regional mutation rates for model comparison.

### Comparison of models with different hyperparameters

Due to the high demand for GPU memory and computing, it is impossible to test the behaviors of all model hyperparameters comprehensively. At the beginning, we set a relatively large search space for hyperparameters and used Ray Tune to run dozens of trials to get more narrowed ranges. We further detailedly investigated the impact of two input-related hyperparameters – ‘local radius’ (length of the local sequence on each side of the focal nucleotide) and ‘distal radius’ (length of the expanded sequence on each side of the focal nucleotide). For each hyperparameter, we set five different values for it and fixed the setting of other hyperparameters. We used a training dataset consisting of 100,000 mutated and 2,000,000 non-mutated A/T sites and a validation dataset consisting of 50,000 mutated and 1,000,000 non-mutated A/T sites. For each setting of hyperparameters, we ran ten trials and used the model with lowest validation loss of each trial for comparison (**Supplementary Fig. 3**).

We also investigated the impact of different training data sizes. We tried four different numbers of A/T sites as training data: 1) 50,000 mutated+1,000,000 non-mutated; 2) 100,000 mutated+2,000,000 non-mutated; 3) 200,000 mutated+4,000,000 non-mutated; and 4) 500,000 mutated+10,000,000 non-mutated. For model evaluation, we used a validation dataset consisting of 50,000 mutated and 1,000,000 non-mutated A/T sites to calculate validation losses for comparison (**Supplementary Fig. 5**).

### Comparison of MuRaL models with previously published models

We considered the following four published models in our comparative analysis:

1. ‘Aggarwala 7-mer’ model^11^: this model estimated 7-mer mutation rates based on intergenic polymorphic sites from 1000 Genomes Project (~11 million variants in the African populations, ~7 million variants in the European populations, and ~6 million variants in the East Asian populations.). The original study provided 7-mer mutation rates for three populations (‘Supplementary Table 7’). We used the averaged mutilation rate among three populations for each 7-mer to generate mutation rates of all bases in human autosomes.
2. ‘Carlson 7-mer’ model^9^: this model used 7-mer mutation rates estimated from 36 million singleton variants from 3560 individuals. Note that some 7-mers didn’t have any observed mutations and thus had mutation rates of zero, which was a limitation of this method. We downloaded the 7-mer mutation rates from ‘Supplementary Data1’ of the study and generated mutation rates of all bases in human autosomes.
3. ‘Carlson 7-mer+features’ model^9^: this model used 7-mer mutation rates of the ‘Carlson 7-mer’ model and 14 genomic features for modeling. We noticed that some sites had zero mutation rates for specific mutation types. In addition, this model did not generate predicted rates for sites within 5 Mb of the start/end of a chromosome because of lacking corresponding recombination rate data. We downloaded the whole genome mutation rate profile of this model from the original study (http://mutation.sph.umich.edu/hg19/) for analysis.
4. ‘Karczewski 3-mer’ model^15^: this model estimated 3-mer mutation rates based on rare variants in gnomAD database. For CpG sites, this model divided the methylation levels into three classes (high, medium and low) and applied separate mutation rates for CpG sites with different methylation levels. We downloaded the 3-mer mutation rates from ‘ Supplementary Dataset 10’ of the study and generated mutation rates of all bases in human autosomes. We used the same methylation data as that described in the study for predicting mutation rates at CpG sites.

When performing comparative analyses between our models and other existing models, we excluded the genomic sites without predictive values in at least one model. In total, 2,390,435,721 bases of the autosome genome were used in comparison. Note that among the four existing models, the ‘Carlson 7-mer+features’ model had strongest data requirements for prediction and its mutation rate profile contains the smallest number of predicted sites. We calculated k-mer and regional mutation rate correlations for the four models using the same method as that for MuRaL models.

The MuRaL models used in comparative analysis were those trained with 500,000 mutated and 10,000,000 non-mutated sites. The number of trainable parameters for each of AT, non-CpG and CpG models was 86,904. The total number of trainable parameters (86,904*3=260,712) was smaller than that of the ‘Carlson 7-mer+features’ model (392,128)^9^.

### Transfer learning

Transfer learning is widely used in deep learning for scenarios in which the prediction tasks are similar. After training MuRaL models with rare variants from gnomAD, we took advantage of published human DNMs to perform transfer learning. For each of the AT and non-CpG models, we compiled a training dataset consisting of 150,000 DNMs and 3,000,000 non-mutated sites and an independent validation dataset consisting of 20,000 DNMs and 400,000 non-mutated sites. For the CpG models, we compiled a training dataset consisting of 50,000 DNMs and 1,000,000 non-mutated sites and a validation dataset consisting of 20,000 DNMs and 400,000 non-mutated sites. We tried two transfer learning strategies: 1) using all pre-trained weights for model initialization and re-training all weights and 2) use all pre-trained weights for model initialization but only re-training the weights of last FC layers of two modules. We chose the results of the first strategy for later comparative analysis, as the second strategy led to poor performance. We also trained *ab initio* models using the same DNM training datasets, with the same hyperparameter setting as that for the rare-variant models. Because we found that the collected DNMs were highly depleted in low-complexity regions and segmental duplications, we excluded DNMs located in these regions from training and evaluating models.

### Applying MuRaL to other species

We used MuRaL to train mutation rate models for three other species: *Macaca mulatta, Drosophila melanogaster* and *Arabidopsis thaliana. M. mulatta* is a widely used primate model organism with similar genome size as that of the human genome. *D. melanogaster* and *A. thaliana* are widely used model organisms but with much smaller genomes (169 Mb and 119 Mb, respectively).

The variants of *M. mulatta* were from a recent study^31^ and downloaded from https://hgdownload.soe.ucsc.edu/gbdb/rheMac10/rhesusSNVs/. This dataset included 853 sequenced genomes and 85.7 million variants. We extracted 19,553,394 singleton variants (requiring AC=1 and AN>=1500) of autosomes for training AT and non-CpG models. For training CpG models, we did downsampling to the total allele count (AN) of 1000 and extracted the variants with AC_down_ ≤ 5 (6,422,014 CpG-related rare variants on autosomes). To identify regions with poor mappability in the *M. mulatta* genome, we downloaded raw reads of three individuals (accession numbers: SRR11999190, SRR11999224 and SRR12070989) and mapped them to the rheMac10 assembly using bwa-mem2 ^49^. The peak read depth for alignments of the three libraries was 127, and we kept genomic sites with read depth within the range of 63-190 (2,620,098,971 bp in autosomes in total) for downstream analyses.

We trained *ab initio* models as well as transfer learning models for *M. mulatta*. For *ab initio* models, we compiled a training dataset consisting of 500,000 mutated and 10,000,000 non-mutated sites, and an independent validation dataset consisting of 50,000 mutated and 1,000,000 non-mutated sites. We used the same hyperparameter setting as that for human *ab initio* models. For transfer learning models, we compiled a training dataset consisting of 150,000 mutated and 3,000,000 non-mutated sites and an independent validation dataset consisting of 50,000 mutated and 1,000,000 non-mutated sites. For each model, ten trials were run and the checkpointed model with lowest validation loss among all trials was used for prediction.

The variant file of *A. thaliana* was downloaded from 1001 Genomes project (https://1001genomes.org/)^50^, which included 12,883,854 polymorphic sites for 1135 inbred lines. The variants of each individual in the VCF file were all homozygotes because of long-term inbreeding and thus the lowest AC is 2. We excluded the poorly mapped genomic regions by using the coverage information from the 1001 Genomes project. We first calculated the average read depth across 1135 lines for each nucleotide and the mode of the rounded average depths across the genome was 21. We retained the positions whose average read depth was within the range of 10-30 (102,069,978 sites in total). For training and validating the AT model, we used singleton variants by requiring AC to be 2 and AN of >= 1000. Because there is a high mutation rate of C>T at both CpG and non-CpG C/G sites, the C>T mutations were depleted in the singleton rare variants. We further compiled a rare variant dataset by requiring AC <= 10 and AN >= 1000 for training and validating non-CpG and CpG models. For each of AT, non-CpG and CpG models, we randomly selected 100,000 rare variants and 2,000,000 non-mutated sites for training, and 10,000 rare variants and 200,000 non-mutated sites for validation. For the CpG model, we randomly selected 50,000 rare variants and 1,000,000 non-mutated sites for training, and 5,000 rare variants and 100,000 non-mutated sites for validation.

The variant dataset of *D. melanogaster* used in our analysis was from Drosophila Genetic Reference Panel (DGRP)^51^, which sequenced 205 inbred lines. The original variant file in VCF format contained 3,837,601 polymorphic sites (excluding sex chromosomes and heterochromatic sequences). Because of being derived from inbred lines, each polymorphic site was homozygous for each individual and original AN and AC tags in the variant file were corresponding to counts of individuals rather than alleles. We extracted 702,864 singleton rare variants by requiring AC to be 1 and AN of >= 100. Then the dataset of rare variants was divided into A/T sites (285,374) and C/G sites (418,713), respectively. Because there is little methylation at CpG sites in the genome of *D. melanogaster*^52^ and the mutation rate of CpG>TpG is not exceptionally high in this species, we did not separate non-CpG and CpG C/G sites for training. For each of the AT and CG models, we randomly selected 100,000 rare variants and 2,000,000 non-mutated sites for training, and 10,000 rare variants and 200,000 non-mutated sites for validation.

The configurations of hyperparameters for MuRaL models of the three species were provided in **Supplementary Tables 9-11**. For each training task, the checkpointed model with lowest validation loss among all trials was used for predicting base-wise mutation rates in the whole genome. The calculation of k-mer and regional mutation rate correlations was the same as that for the human data.

### Mutation rates around human genes

We downloaded human gene annotations from GENCODE^53^ (v37) and extracted the longest transcript for each protein-coding gene as its representative transcript. DeepTools^54^ (v3.5.1) was used for plotting meta-gene profiles and heatmaps for mutation rates around gene regions, as well as doing K-means clustering. We tried K-means clustering with multiple K values. Because CpG-related variants were depleted in observed rare variants, we only considered A/T and non-CpG C/G sites for this analysis. ClusterProfiler^55^ (v4.4.4) was used to perform GO enrichment analysis and enriched ‘biological process’ GO terms were used for visualization.

### Depletion rank (DR)

We calculated the DR scores using the same method as that in Halldorsson et al.^33^. For each of sliding 500-bp windows with a 50-bp step across the genome, we computed the observed number of all gnomAD variants (O) and the expected number of variants (E). The expected number of variants (E) was calculated by summing the scaled MuRaL mutation rates in the given window and multiplying the summed rate by a correction factor. The correction factor was the ratio between the observed number of gnomAD variants and the summed mutation rate in a set of presumably neutral sites from a previous study^56^. To account for read coverage heterogeneity, we also used the pre-computed coverage correction model (0.48449*log10(coverage)+0.22009) from gnomAD^15^ (https://github.com/broadinstitute/gnomad_lof) to adjust the mutation rates before summing. With obtained O and E values, we then calculated the metric (O-E)/✓ E for all windows. The window with the i-th lowest (O-E)/✓E was assigned a DR score of 100*(i-0.5)/n, where n is the total number of windows. Due to the saturation issue of CpG variants, we only considered A/T and non-CpG C/G sites for this analysis. The ClinVar variants were downloaded from https://ftp.ncbi.nlm.nih.gov/pub/clinvar/vcf_GRCh37/clinvar_20220730.vcf.gz, and only SNVs were used in our analysis.

## Data availability

All the analyses in this study were based on published data. The predicted mutation rate maps for genomes of human, *M. mulatta, A. thaliana* and *D. melanogaster* are available at the ScienceDB repository: https://www.doi.org/10.11922/sciencedb.01173.

## Code availability

Source code of the MuRaL package is available at https://github.com/CaiLiLab/MuRaL. The MuRaL version (v1.0.0) used for this publication is also available at Zenodo (https://doi.org/10.5281/zenodo.6989025)

## Acknowledgements

We thank Xionglei He, Xia Shen and Nicholas Luscombe for giving insightful comments on the manuscript. We thank all lab members for discussion and help throughout this project. This work was supported by National Natural Science Foundation of China (32070593), Guangdong Basic and Applied Basic Research Foundation (2022A1515010888), and Science and Technology Planning Project of Guangzhou (202102020816).

## Author contributions

C.L. designed and supervised the project. C.L. developed the MuRaL framework, with input from Y.F. and S.D. for detailed evaluation. Y.F. and S.D. performed comparative analyses and generated mutation rate profiles. C.L, Y.F. and S.D. wrote the manuscript.

