## Supplementary information for "A generalizable deep learning framework for inferring fine-scale germline mutation rate maps"

**This document contains Supplementary Figures 1-23 and Supplementary Tables 1-11.**

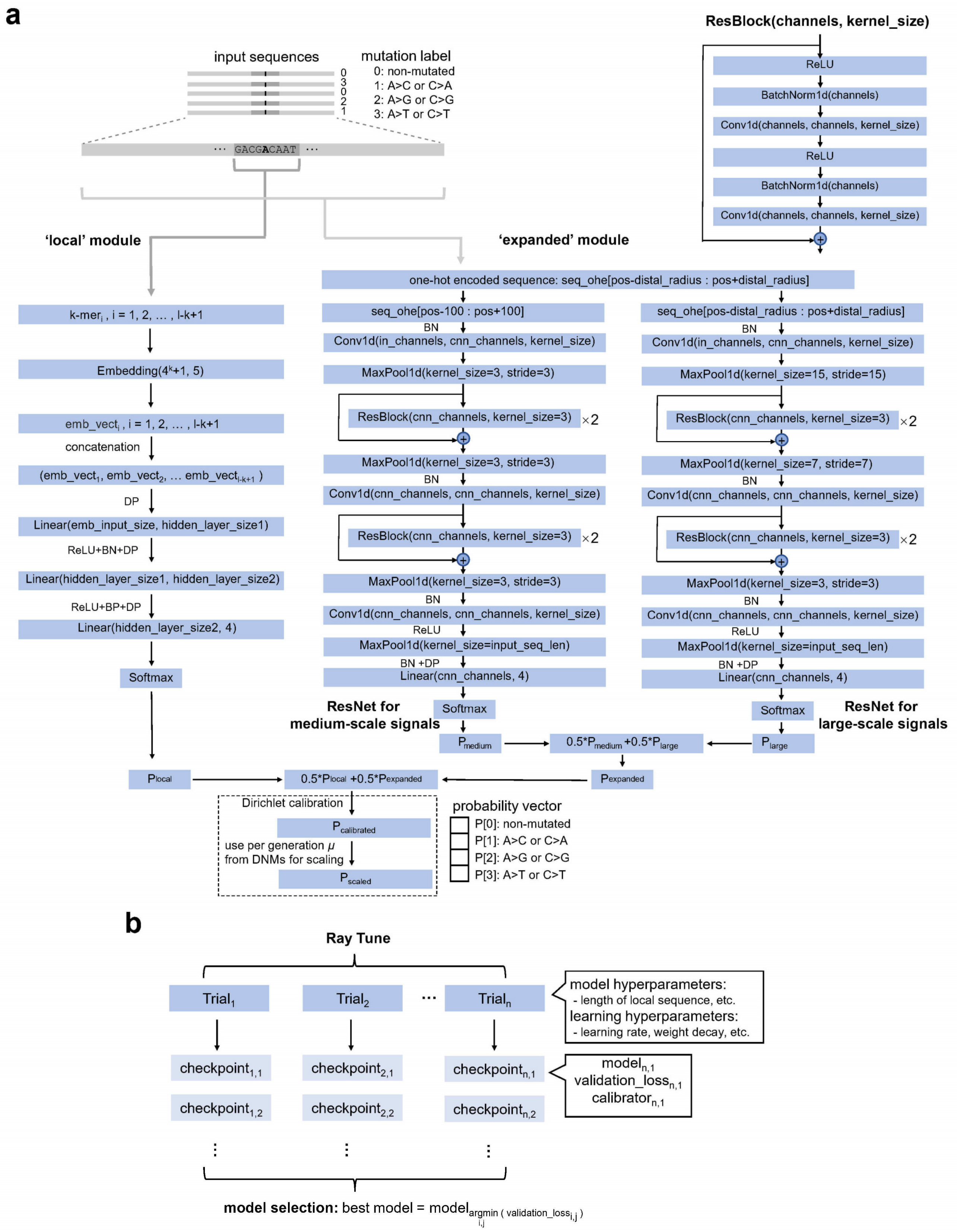


**Supplementary Figure 1 Illustration for MuRaL layers and model training.** (**a**)The MuRaL model has a 'local' module and a 'expanded' module. In the 'local' module, the input sequence of the focal nucleotide (i.e., 'A' in the figure) is firstly split into overlapping k-mers, which are then mapped into multi-dimensional vectors by the embedding layer. The resulting multi-dimensional vectors are concatenated to form the input for subsequent two hidden FC layers and one output FC layers. For each hidden layer, ReLU activation function is used with the output of the FC layer, followed by batch normalization (BN) and dropout (DP) layers. The output of the 'local' module is a probability distribution generated by the softmax function over four predicted classes - being non-mutated or mutated to another three possible nucleotides. In the 'expanded' module, the input sequence of an expanded region is first one-hot encoded. The one-hot encoded matrix is considered as one-dimensional data with four channels and passed to a Residual Network (ResNet) component, which has two ResNets for learning middle-scale and large-scale sequence signals, respectively. Each ResNet has two sets of stacked residual blocks that are placed between three separate one-dimensional CNN layers. The design of a single residual block is provided in the box on the top right. Three maxpooling layers are used to gradually reduce the sequence size. An additional FC layer and the softmax function following the ResNet component generate a probability distribution over four predicted classes, like that in the 'local' module. The probabilities of 'medium-scale' and 'large-scale' ResNets are combined with equal weights to form the probabilities of the 'expanded' module. The probabilities of 'local' and 'expanded' modules are combined using equal weights to form a vector of combined probabilities. The combined probabilities can be better calibrated by the Dirichlet calibration and further scaled to per-generation mutation rates with known average genome-wide mutation rate estimates. Note that the calibration and scaling steps in the dashed box do not take part in neural network training. (**b**) Illustration for model training with Ray Tune and model selection. For training a model, a certain number of trials are run with a specific set of hyperparameters, using the same training and validation datasets. For each trial, the trained model parameters and the calibrator are saved at each checkpoint (e.g., at the end of each training epoch). After training, the model with lowest validation loss is considered as the best model, which can be used for prediction tasks.

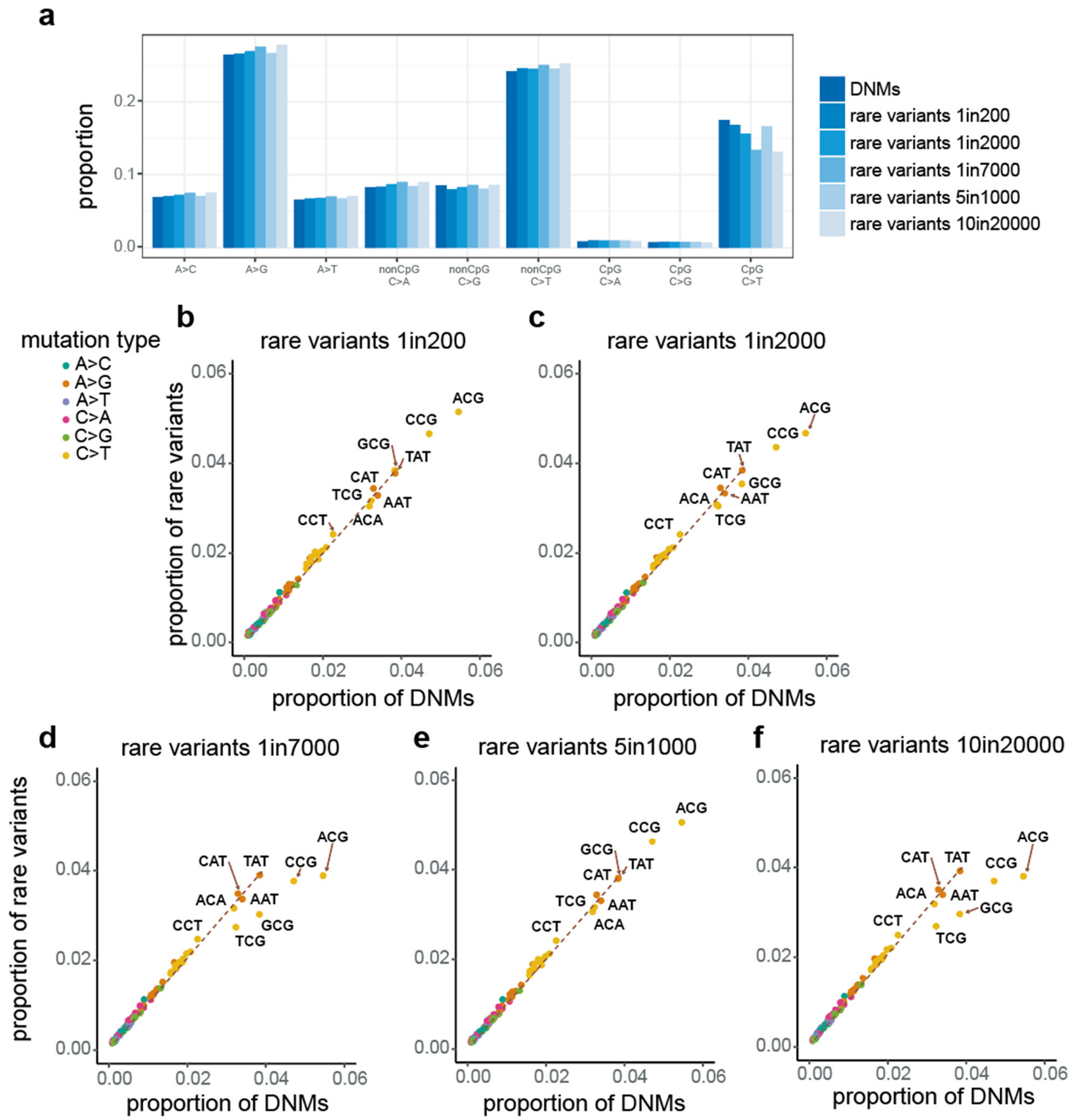


**Supplementary Figure 2 Comparison of 1-mer and 3-mer mutation spectra of rare variants from different downsampled datasets with those of DNMs.** (**a**) Comparison of proportions of 1-mer mutation types. (**b-f**) Comparison of proportions of 3-mer mutation subtypes of rare variants and that of DNMs. The dotted lines in panels **b-f** were fitted with all 3-mer mutation subtypes except four CpG>TpG subtypes (A[C>T]G, C[C>T]G, G[C>T]G and T[C>T]G).

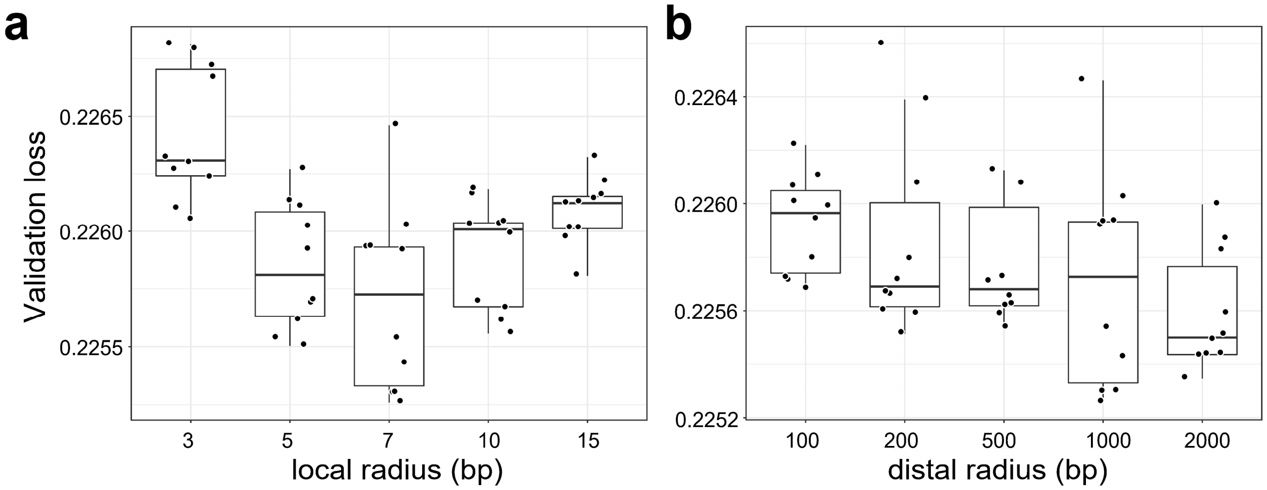


**Supplementary Figure 3 Validation losses for models with different lengths of 'local' and 'expanded' regions.** (**a**) Validation losses for models with different local radiuses. Here local radius is the number of nucleotides on each side of the focal nucleotide of the 'local' region. The shown boxplots were based on results of AT models trained with 100,000 mutated and 2,000,000 non-mutated. For each model, the lowest loss (mean cross-entropy loss) in each of ten trials was used to generate the boxplots. (**b**) Validation losses for models with different distal radiuses. Distal radius is the number of nucleotides on each side of the focal nucleotide of the 'expanded' region. The boxplots were generated in the same manner as that for panel **a**.

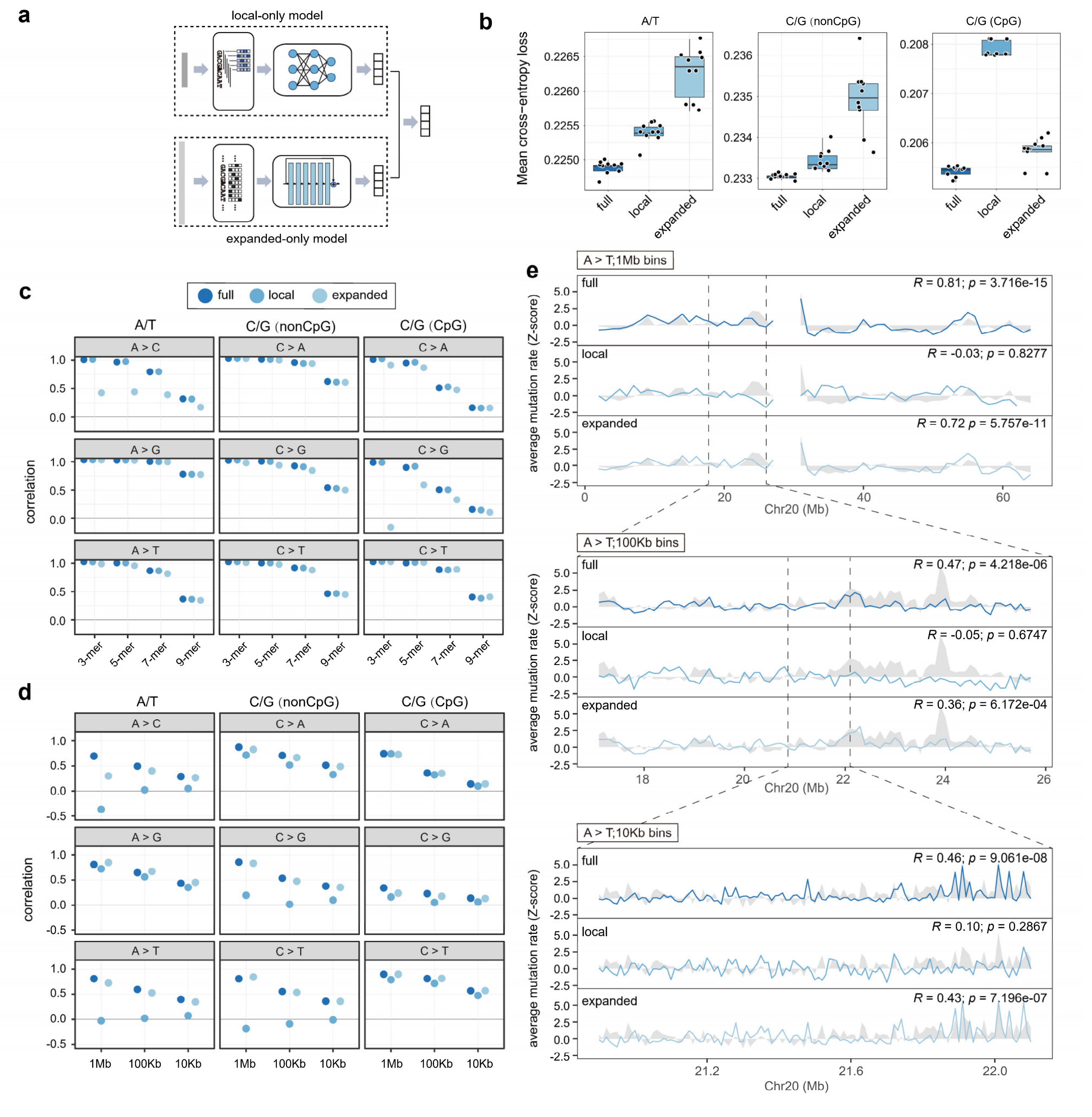


**Supplementary Figure 4 Two modules of MuRaL learn different mutability signals.** (**a**) Illustration of 'local-only' and 'expanded-only' models. Diagram elements have same meanings as that in Fig. 1. (**b**) Average validation losses for MuRaL models of three architectures (full, 'local-only' and 'expanded-only'). Separate models were trained for A/T sites, non-CpG C/G sites and CpG sites, respectively. For each model, the lowest loss for each of ten trials was used to do the boxplots. (**c**) 3-, 5-, 7- and 9-mer mutation rate correlations for different mutation types, based on predicted single-nucleotide mutation rates on human chromosome 20 (Chr20) by models of three different architectures. For each architecture in panel **b**, the best trial with lowest validation loss was used for prediction. The mutations for calculating observed mutation rates were '10in20000' rare variants for AT and non-CpG models, and '5in1000' rare variants for CpG models. (**d**) Regional mutation rate correlations with bin sizes of 1Mb, 100Kb and 10Kb on Chr20 for different mutation types. Used observed mutations were the same as that for panel **c**. (**e**) An example showing regional A>T mutation rate correlations at different scales on Chr20 for three models, with grey shades indicating observed rates and colored lines for predicted rates. Used models and observed mutations were the same as that for panel **d**. As predicted and observed mutation rates had different magnitudes, z-score normalization was applied for visualization. Mutation rates at centromeric regions were not available. Pearson's coefficients and p-values for shown regions are provided at the upper right corners.

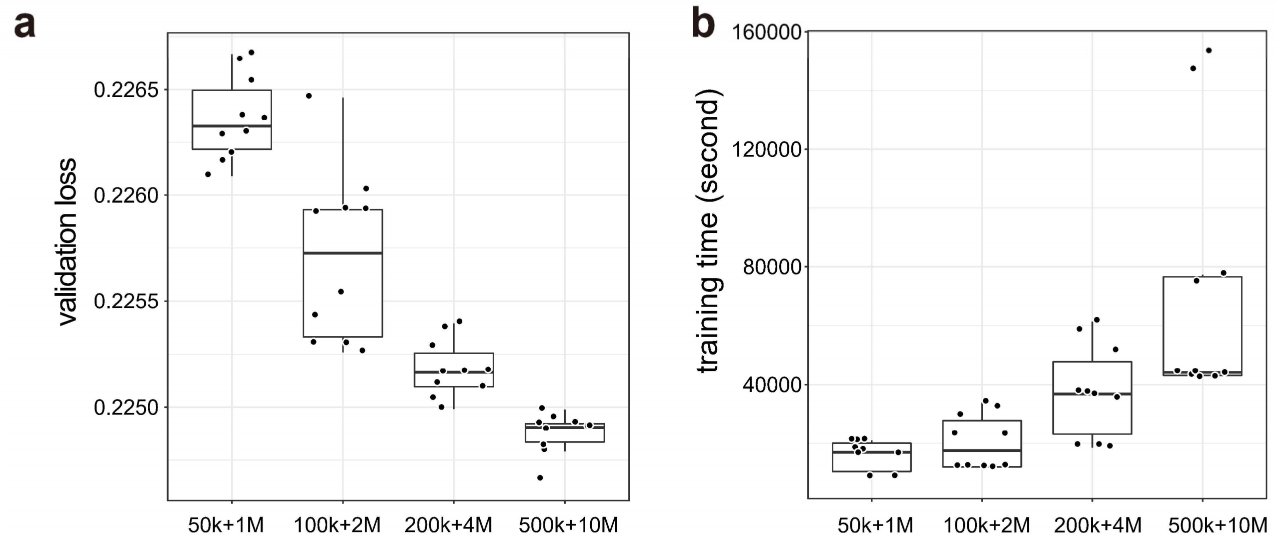


**Supplementary Figure 5 Comparison of models trained with different training data sizes.** (**a**) Validation losses for AT models with different training data sizes. For example, '50K+1M' means 50,000 mutated and 1,000,000 non-mutated A/T sites for training. Ten trials were run for each data size. The lowest loss (mean cross-entropy loss) in each of ten trials was used to generate the boxplots. (**b**) Training time for models with different training data sizes. Each point indicates the training time of one specific trial in panel **a**. Training jobs were run on a server with a GeForce RTX 3090 GPU and 256GB RAM.

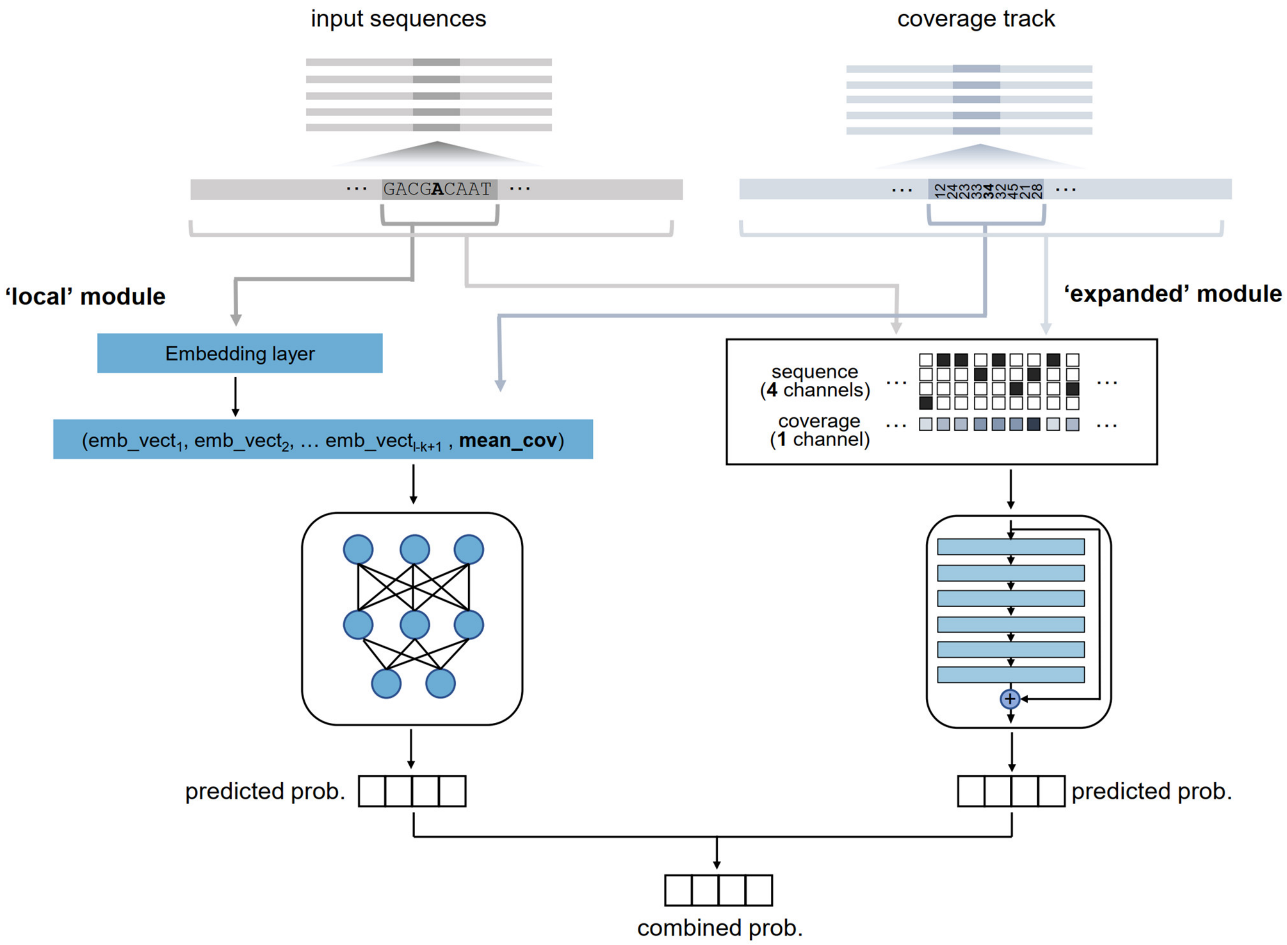


**Supplementary Figure 6 Schematic for incorporating the coverage track into the MuRaL model.** We extended the MuRaL model to incorporate the base-wise coverage information (e.g., in bigWig format). In the 'local' module, we calculated the mean coverage of the local sequence of a focal nucleotide, and added it as an additional element to the concatenated vector of embeddings for the local sequence. In the 'expanded' module, we extracted a coverage vector of the same length as that for the expanded sequence, and merged it with the one-hot encoded matrix of the expanded sequence to form a five-channel input for subsequent convolutional networks. Our design can also easily incorporate other genomic tracks to extend the MuRaL model.

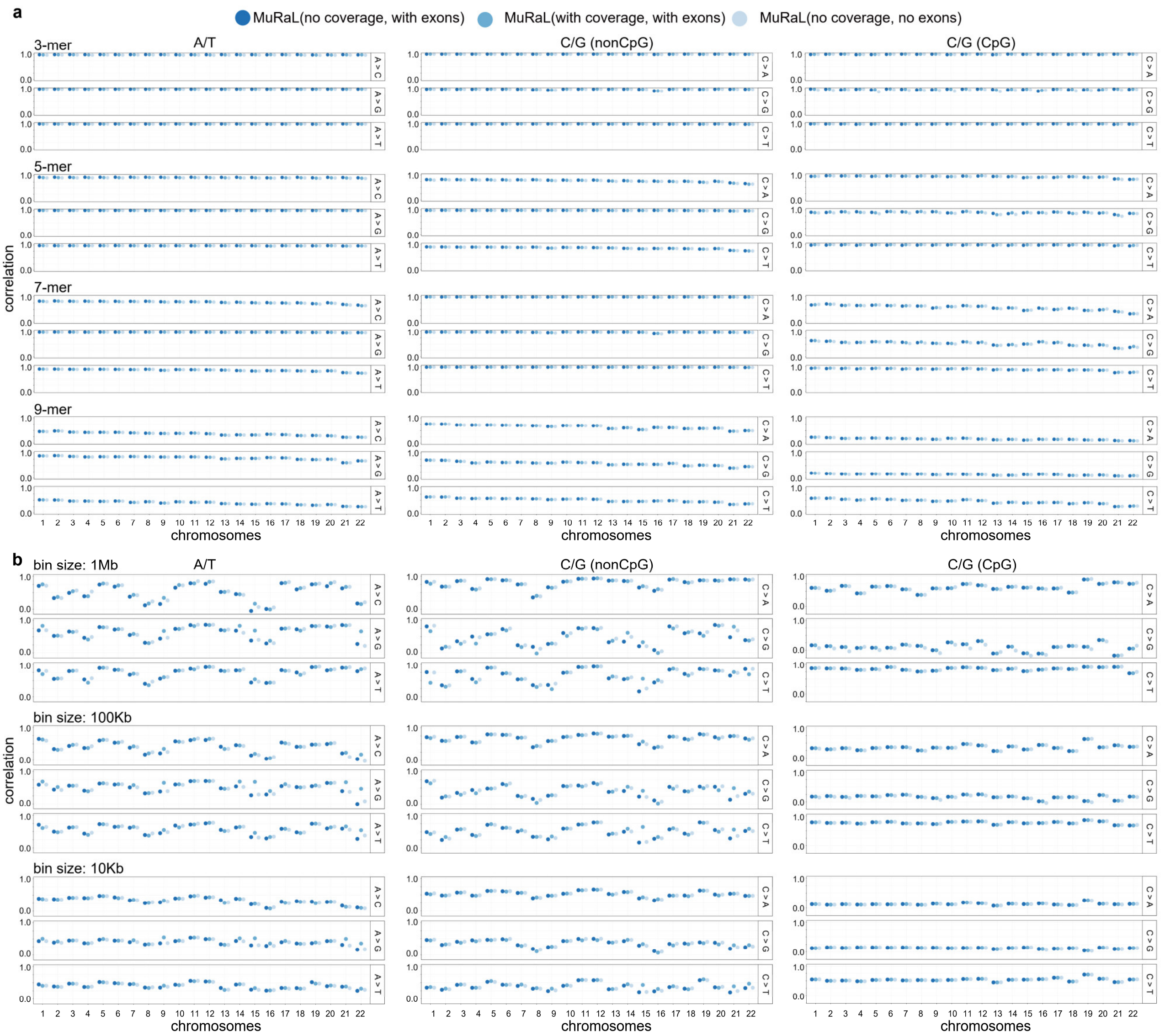


**Supplementary Figure 7 Comparison of MuRaL models trained with different input data.** (**a**) 3-, 5-, 7- and 9-mer mutation rate correlations for different mutation types in each autosome, with colors indicating whether coverage information was used for training. Separate models were trained for A/T sites, non-CpG C/G sites and CpG sites, respectively. The mutations for calculating observed mutation rates were ‘10in20000’ rare variants for AT and non-CpG models, and ‘5in1000’ rare variants for CpG models. In this analysis we considered well-covered regions within mean read coverage ranging of from 15 to 45, which generally had high mappabilities. (**b**) Regional mutation rate correlations with bin sizes of 1Mb, 100Kb and 10Kb for different mutation types in each autosome. The models without coverage tracks but with exonic variants in training data (dark blue) were used for most downstream analyses.

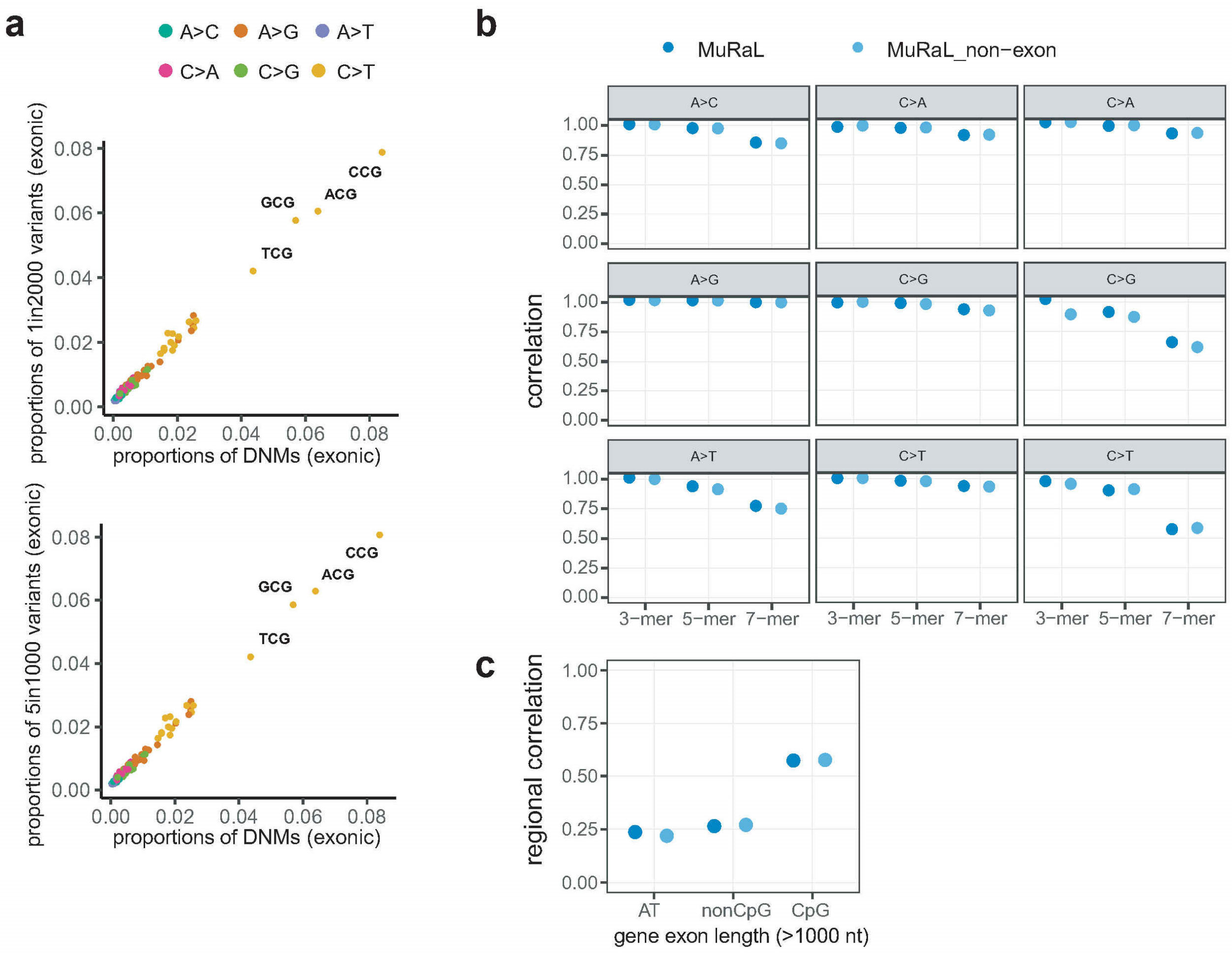


**Supplementary Figure 8 Comparison of models trained with or without exonic rare variants.** (**a**) Comparison of proportions of 3-mer mutation subtypes between exonic rare variants (upper, '1in2000'; lower, '5in1000') and that of exonic DNMs. (**b**) 3-, 5- and 7-mer mutation rate correlations for different mutation types in exonic regions, based on predicted single-nucleotide mutation rates on all autosomes by two models (with or without exonic rare variants for model training). The mutations for calculating observed mutation rates were '10in20000' rare variants for AT and non-CpG models, and '5in1000' rare variants for CpG models. (**c**) Regional mutation rate correlations of two models for different mutation types. Here the average observed and predicted mutation rates were calculated for exonic regions of each transcript. Only the transcripts with a total exon length of >1000 nt were considered. Because usually few mutations were observed in a single transcript, we aggregated mutation types for A/T, non-CpG C/G and CpG sites, respectively, for this analysis.

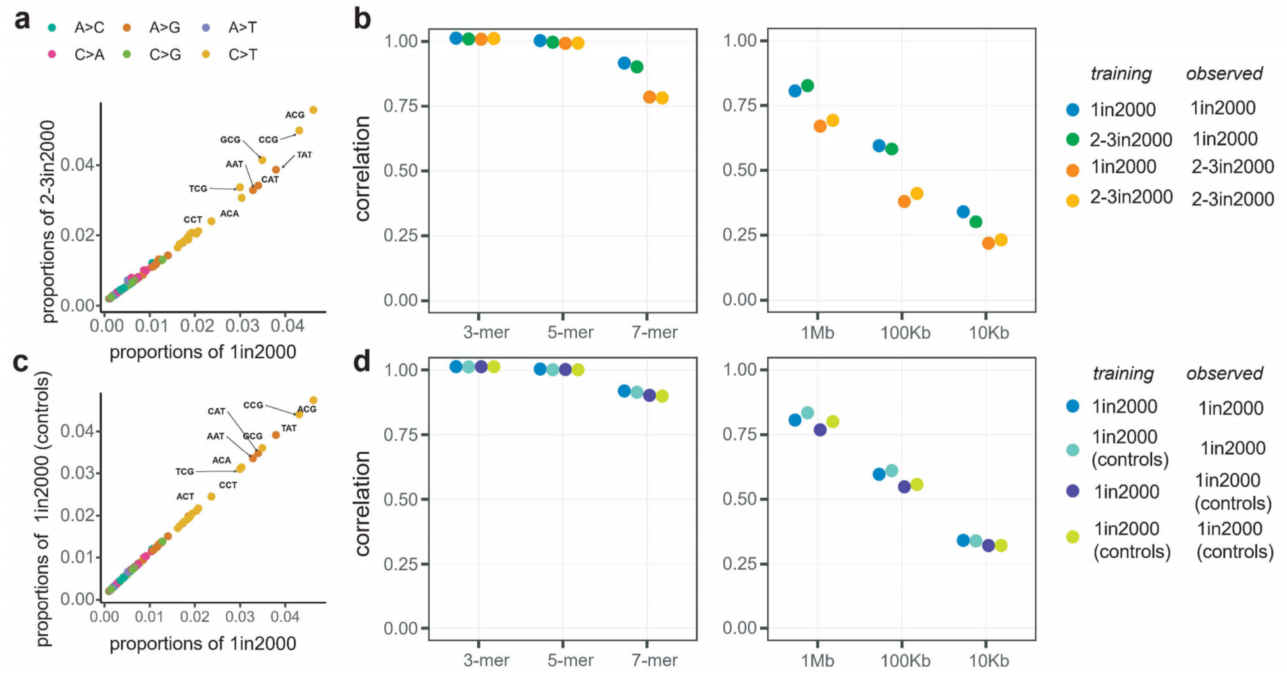


**Supplementary Figure 9 Training models using rare variants after excluding singletons or using rare variants from only healthy controls in gnomAD.** (**a**) Comparison of proportions of 3-mer mutation subtypes of ‘1in2000’ variants and that of ‘2-3in2000’ variants. Here ‘2-3in2000’ means the variants occurring 2 or 3 times after downsampling the total allele count to 2000. The higher proportions of CpG>TpG mutations in ‘2-3in2000’ is expected, given their high mutation rates. (**b**) K-mer (left) and regional (right) mutation rate correlations for AT models trained with ‘1in2000’ or ‘2-3in2000’ data, based on predicted single-nucleotide mutation rates of Chr20. Two sets of rare variants, ‘1in2000’ or ‘2-3in2000’ variants on Chr20, were used for calculating observed mutation rates. The patterns suggest that both models have similar performance. (**c-d**) Similar to the analyses in panels **a-b**, except that the downsampled ‘1in2000’ data from only healthy control individuals in gnomAD was used for comparison.

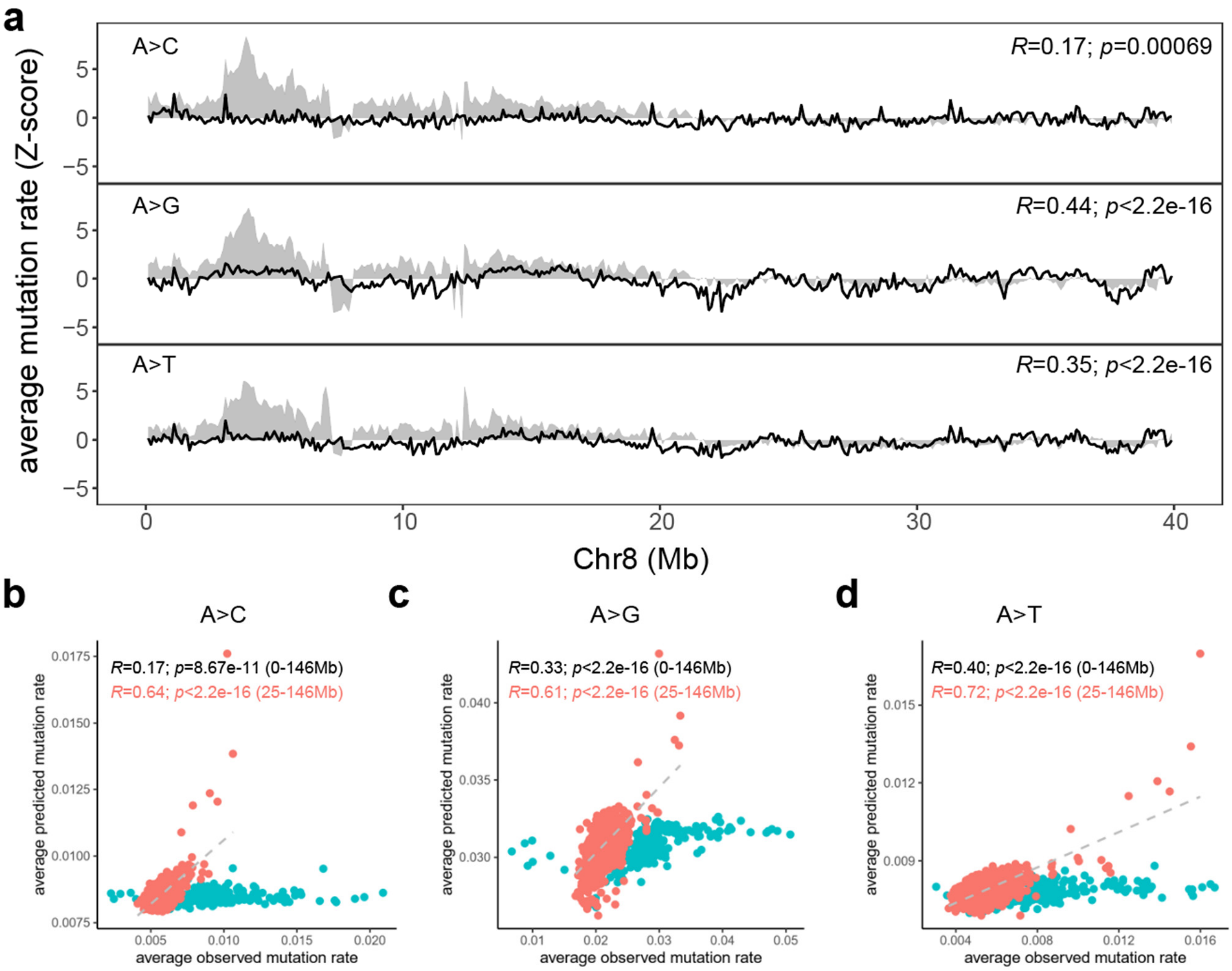


**Supplementary Figure 10 Poor regional correlations on human Chr8.** (**a**) The observed (grey shades) and predicted (black lines) regional mutation rates of 100Kb bins for three mutation types (A>C, A>G and A>T) are shown for the 0-40Mb of human Chr8. Pearson correlation coefficients and p-values for shown regions are provided at the upper right corners. The mutations for calculating observed mutation rates were '10in20000' rare variants. (**b-d**) Scatterplots for the observed and predicted regional mutation rates of 100Kb bins on human Chr8 for three mutation types, with pink dots depicting the bins from 0-25Mb and red dots for bins from 25-146Mb. Pearson correlation coefficients and p-values for all dots (0-146Mb) and red dots (25-146Mb) are shown at the upper left corners.

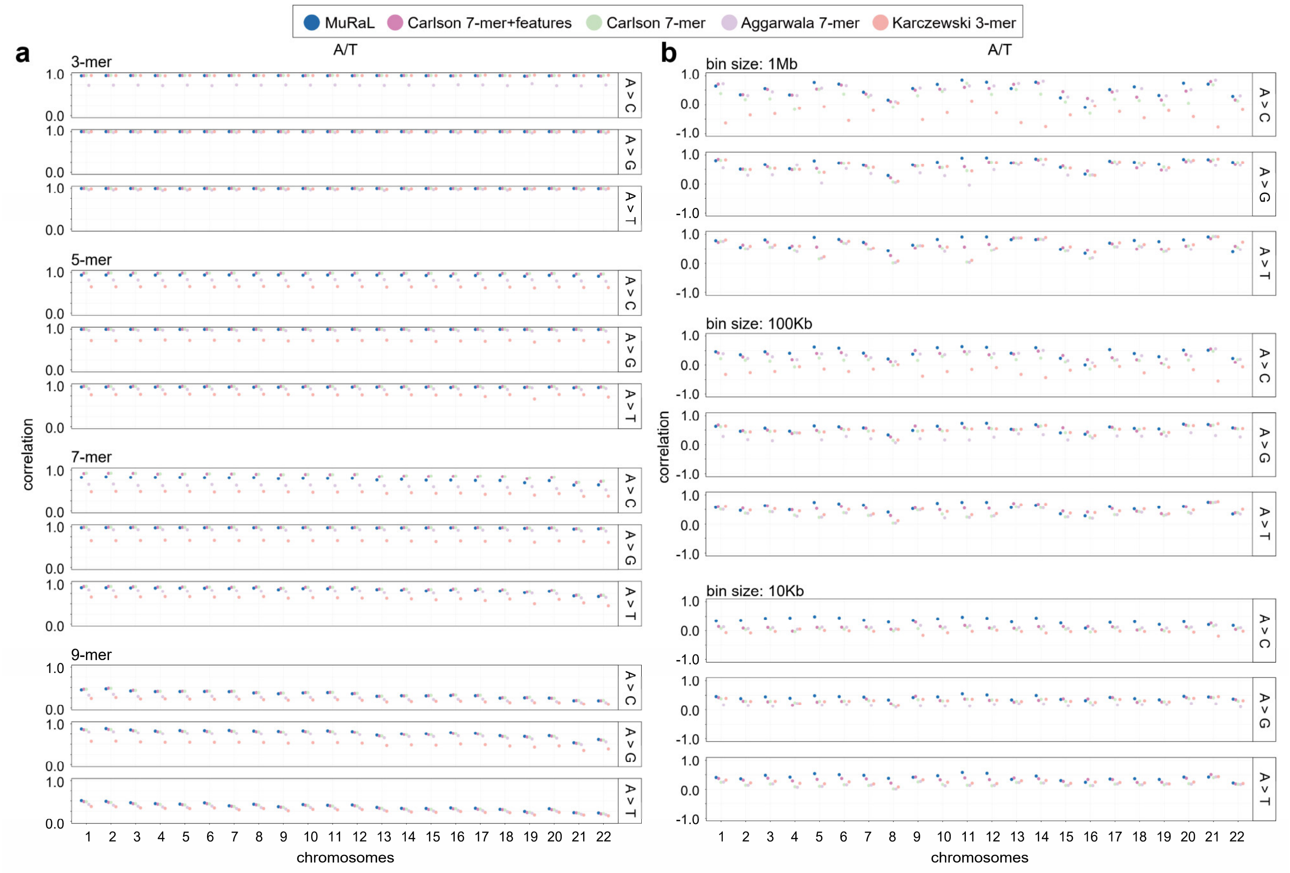


**Supplementary Figure 11 Comparison of k-mer and regional mutation rate correlations between MuRaL and previous models at A/T sites.** (**a**) 3-, 5-, 7- and 9-mer mutation rate correlations for different mutation types in each autosome, with colors indicating different models. The mutations for calculating observed mutation rates were '10in20000' rare variants. (**b**) Regional mutation rate correlations with bin sizes of 1Mb, 100Kb and 10Kb for different mutation types in each autosome.

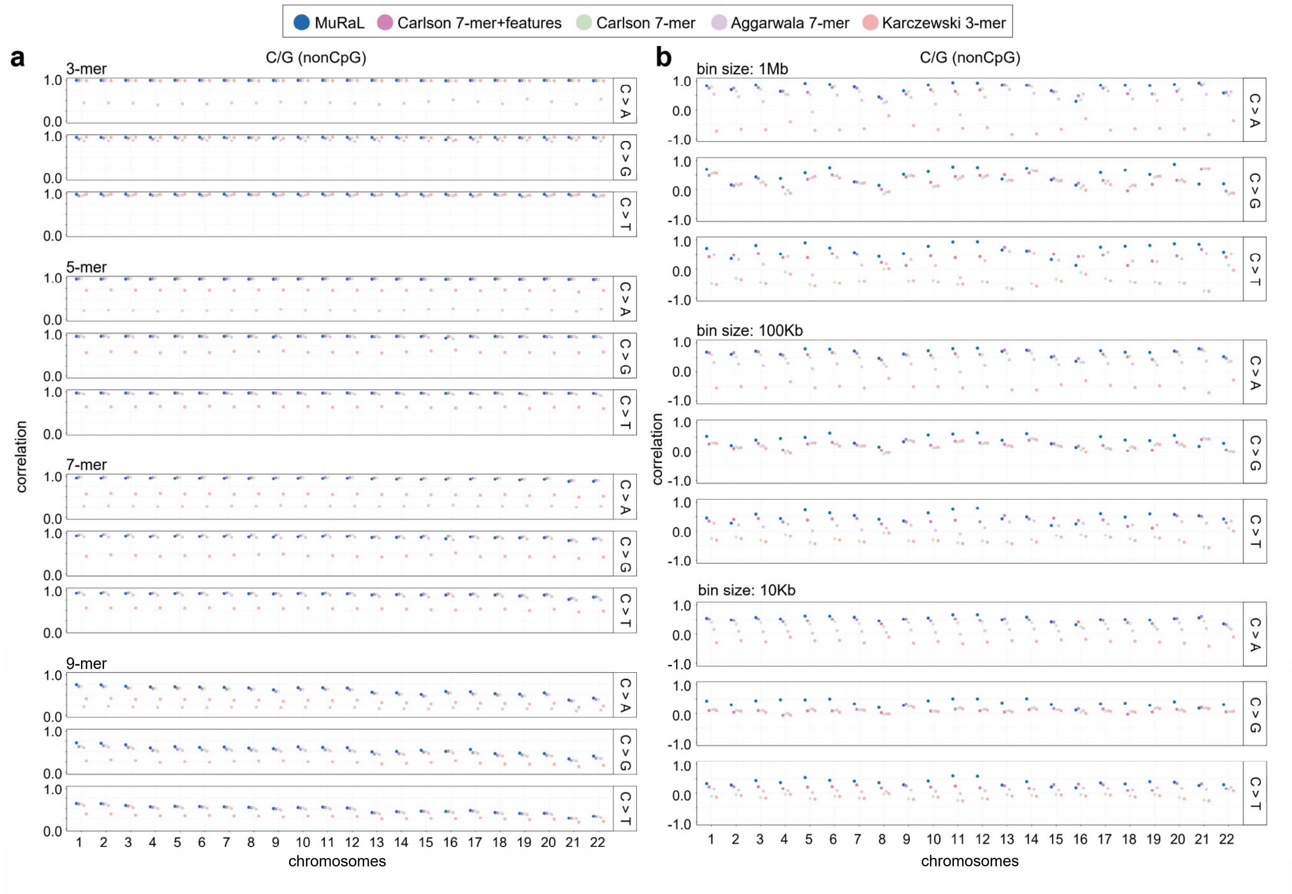


**Supplementary Figure 12 Comparison of k-mer and regional mutation rate correlations between MuRaL and previous models at non-CpG C/G sites.** (**a**) 3-, 5-, 7- and 9-mer mutation rate correlations for different mutation types in each autosome, with colors indicating different models. The mutations for calculating observed mutation rates were '10in20000' rare variants. (**b**) Regional mutation rate correlations with bin sizes of 1Mb, 100Kb and 10Kb for different mutation types in each autosome.

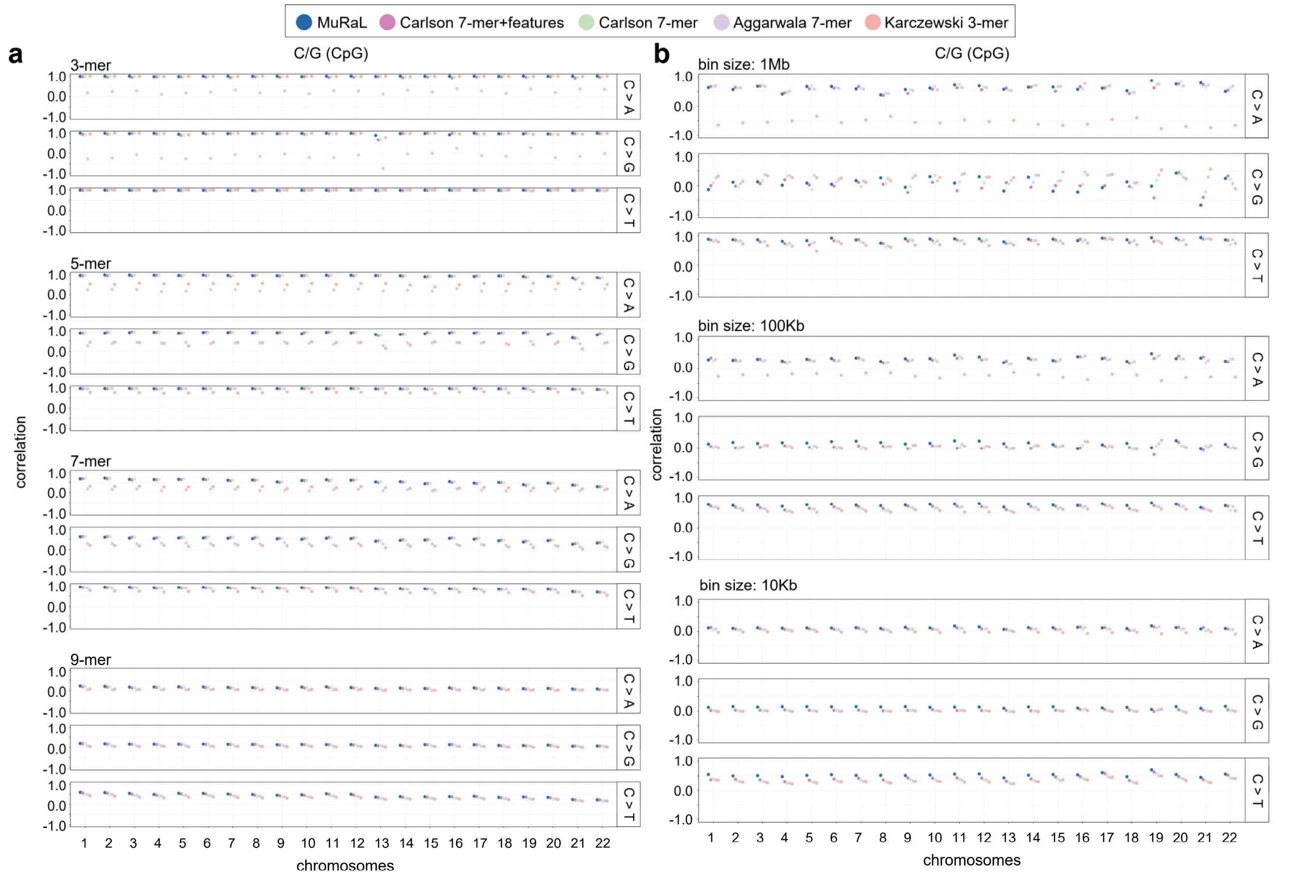


**Supplementary Figure 13 Comparison of k-mer and regional mutation rate correlations between MuRaL and previous models at CpG C/G sites.** (**a**) 3-, 5-, 7- and 9-mer mutation rate correlations for different mutation types in each autosome, with colors indicating different models. The mutations for calculating observed mutation rates were ‘5in1000’ rare variants (**b**) Regional mutation rate correlations with bin sizes of 1Mb, 100Kb and 10Kb for different mutation types in each autosome.

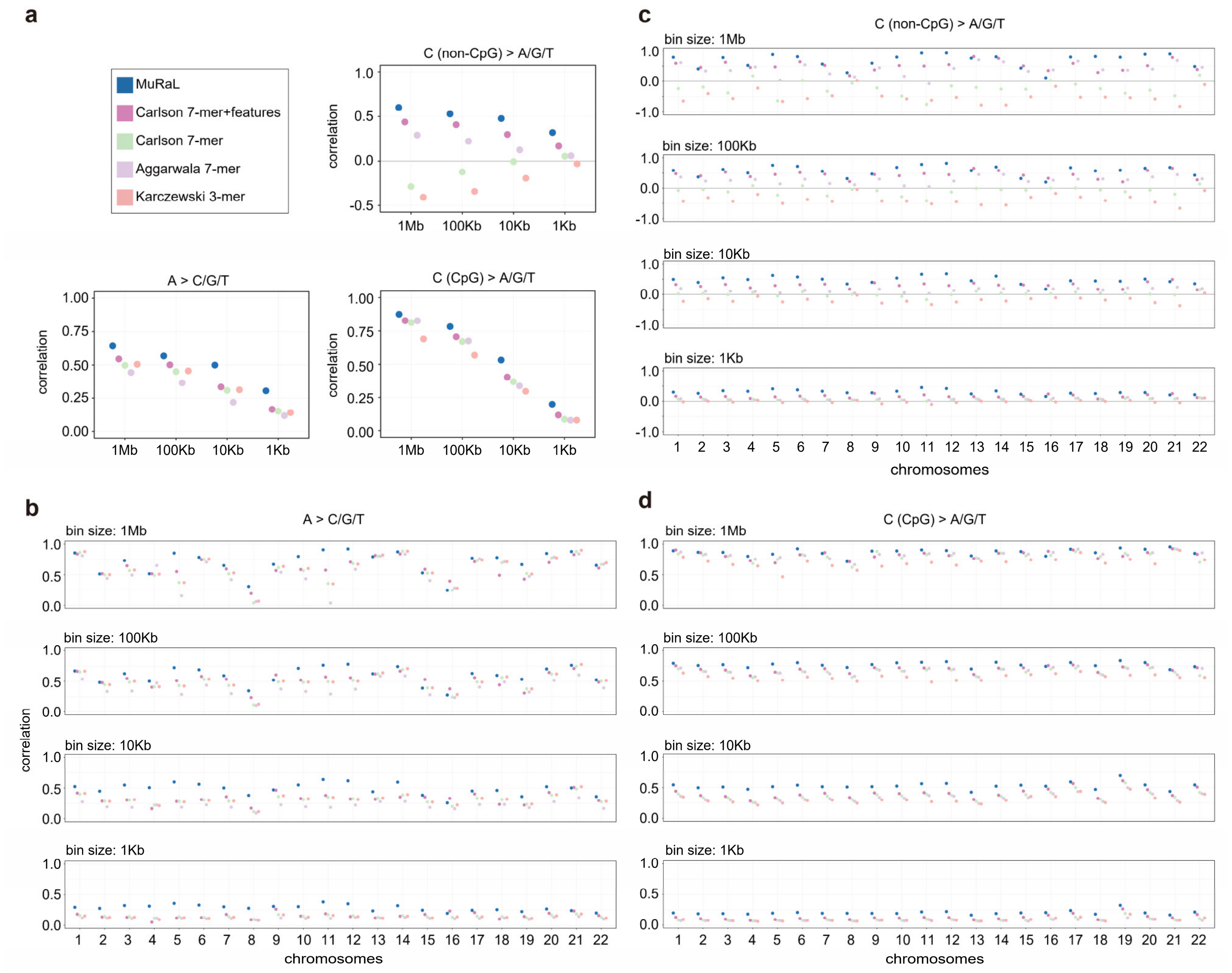


**Supplementary Figure 14 Comparison of regional mutation rate correlations between MuRaL and previous models after aggregating mutation types.** (**a**) Regional mutation rate correlations with bin sizes of 1Mb, 100Kb, 10Kb and 1Kb on the autosomal genome, aggregating mutation types associated with the same reference base. The mutations for calculating observed mutation rates were ‘10in20000’ rare variants for AT and non-CpG models, and ‘5in1000’ rare variants for CpG models. (**b**) Regional mutation rate correlations with bin sizes of 1Mb, 100Kb,10Kb and 1Kb for mutations at A/T sites in each autosome, aggregating A>C, A>G and A>T mutations. (**c**) Regional mutation rate correlations with bin sizes of 1Mb, 100Kb,10Kb and 1Kb for mutations at non-CpG C/G sites in each autosome, aggregating C>A, C>G and C>T mutations. (**d**) Regional mutation rate correlations with bin sizes of 1Mb, 100Kb,10Kb and 1Kb for mutations at CpG C/G sites in each autosome, aggregating C>A, C>G and C>T mutations.

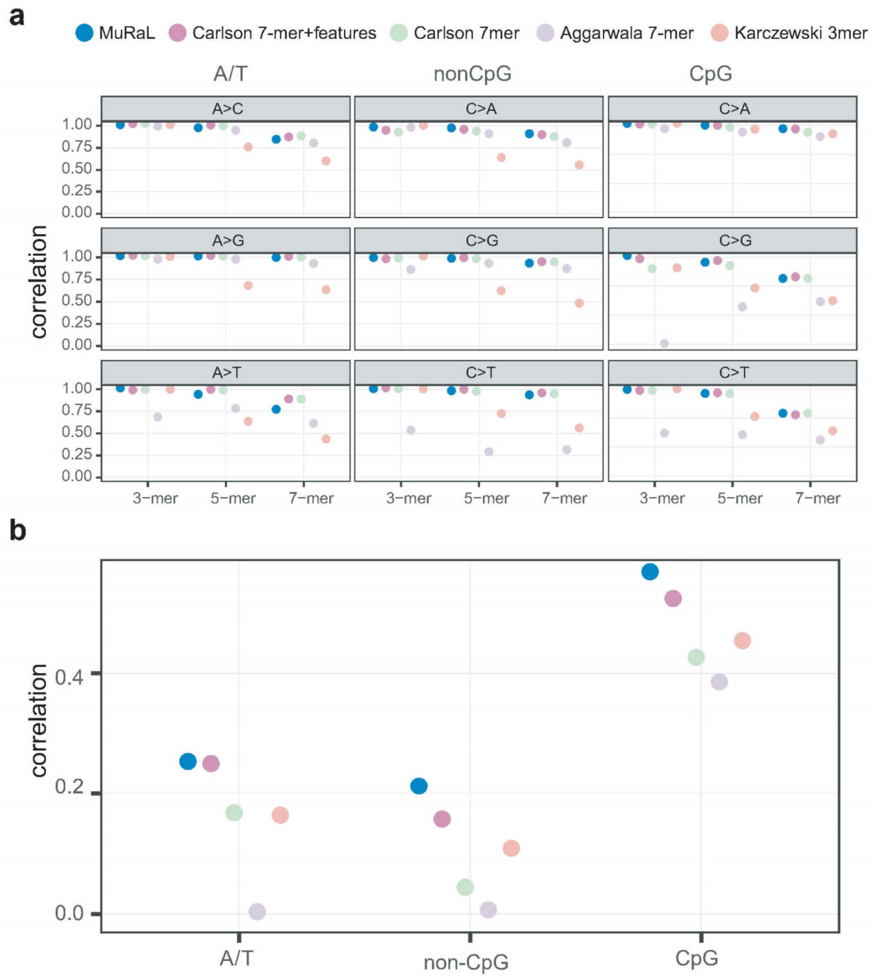


**Supplementary Figure 15 Comparison of k-mer/regional mutation rate correlations between MuRaL and previous models in exonic regions.** (**a**) 3-, 5- and 7-mer mutation rate correlations for different mutation types in exonic regions, based on predicted single-nucleotide mutation rates on all autosomes by MuRaL and previous models. The mutations for calculating observed mutation rates were '10in20000' rare variants for AT and non-CpG models, and '5in1000' rare variants for CpG models. (**b**) Regional mutation rate correlations of five models for different mutation types. Here the average observed and predicted mutation rates were calculated for exonic regions of each transcript. Only the transcripts with a total exon length of >1000 nt were considered. Because of few mutations observed in a single transcript, we aggregated mutation types for A/T, non-CpG C/G and CpG sites, respectively, for analysis.

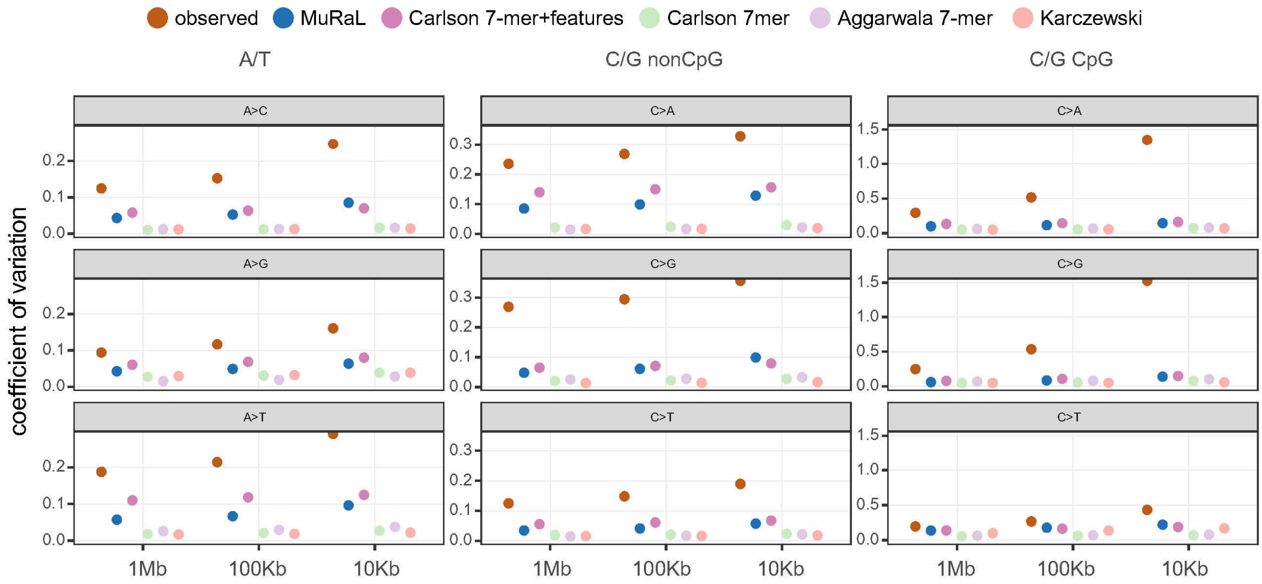


**Supplementary Figure 16 Coefficients of variation for regional mutation rates of different models.** The mutations for calculating observed mutation rates were ‘10in20000’ rare variants for AT and non-CpG models, and ‘5in1000’ rare variants for CpG models.

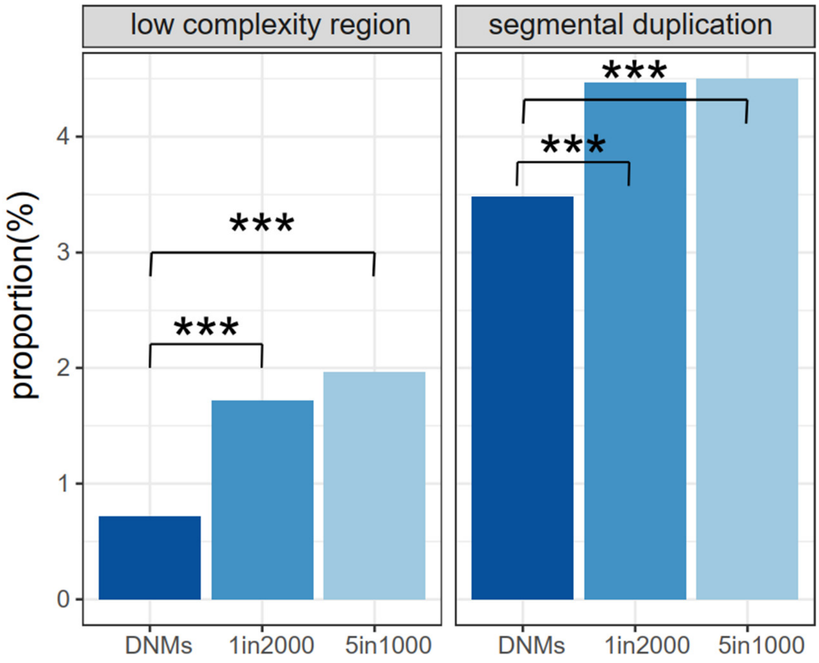


**Supplementary Figure 17 Depletion of mutations in low-complexity regions and segmental duplications in DNM data, in comparison with rare variant datasets ('1in2000' and '5in1000') derived from gnomAD data.** ***, p-value < 2.2e-16 for two-sided Fisher's exact tests.

‘’

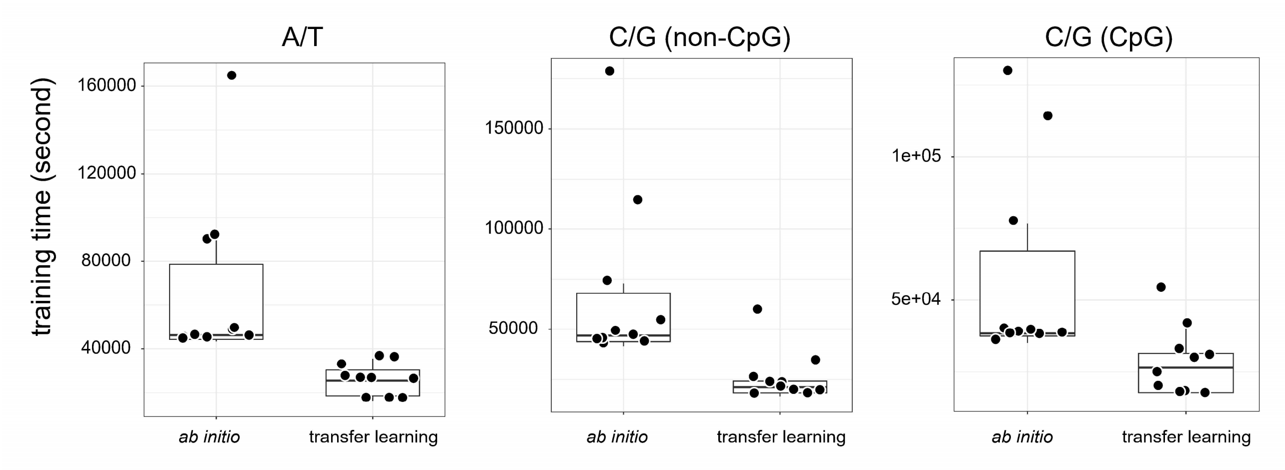


**Supplementary Figure 18 Comparison of training time between *ab initio* and transfer learning models for *M. mulatta*.** Ten trials were run for each model. Each point indicates the training time of one specific trial.

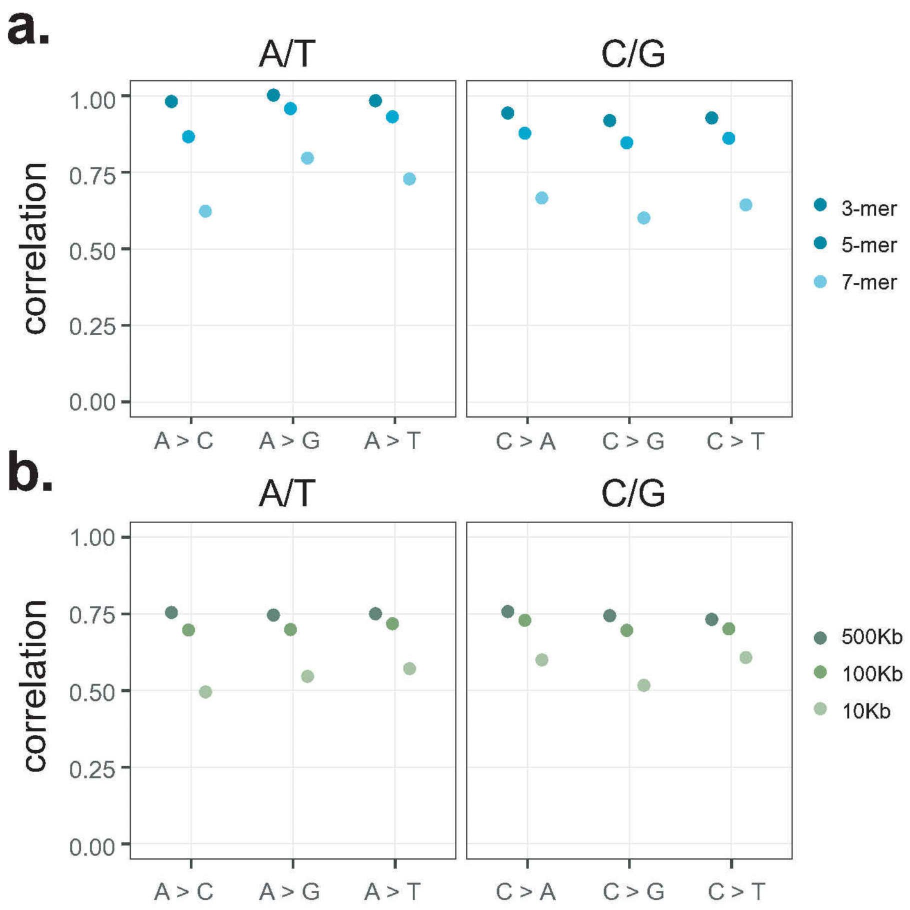


**Supplementary Figure 19 K-mer and regional mutation rate correlations for models of *D. melanogaster.*** (**a**) 3-, 5-, and 7-mer mutation rate correlations for different mutation types, based on predicted single-nucleotide mutation rates for the genome of *D. melanogaster*. Singleton variants of *D. melanogaster* were used for calculating observed mutation rates. Separate models were trained for A/T sites and C/G sites, respectively. (**b**) Regional mutation rate correlations with bin sizes of 500Kb, 100Kb and 10Kb for the *D. melanogaster* genome. Rare variants for calculating observed mutation rates were the same as that for panel **a**.

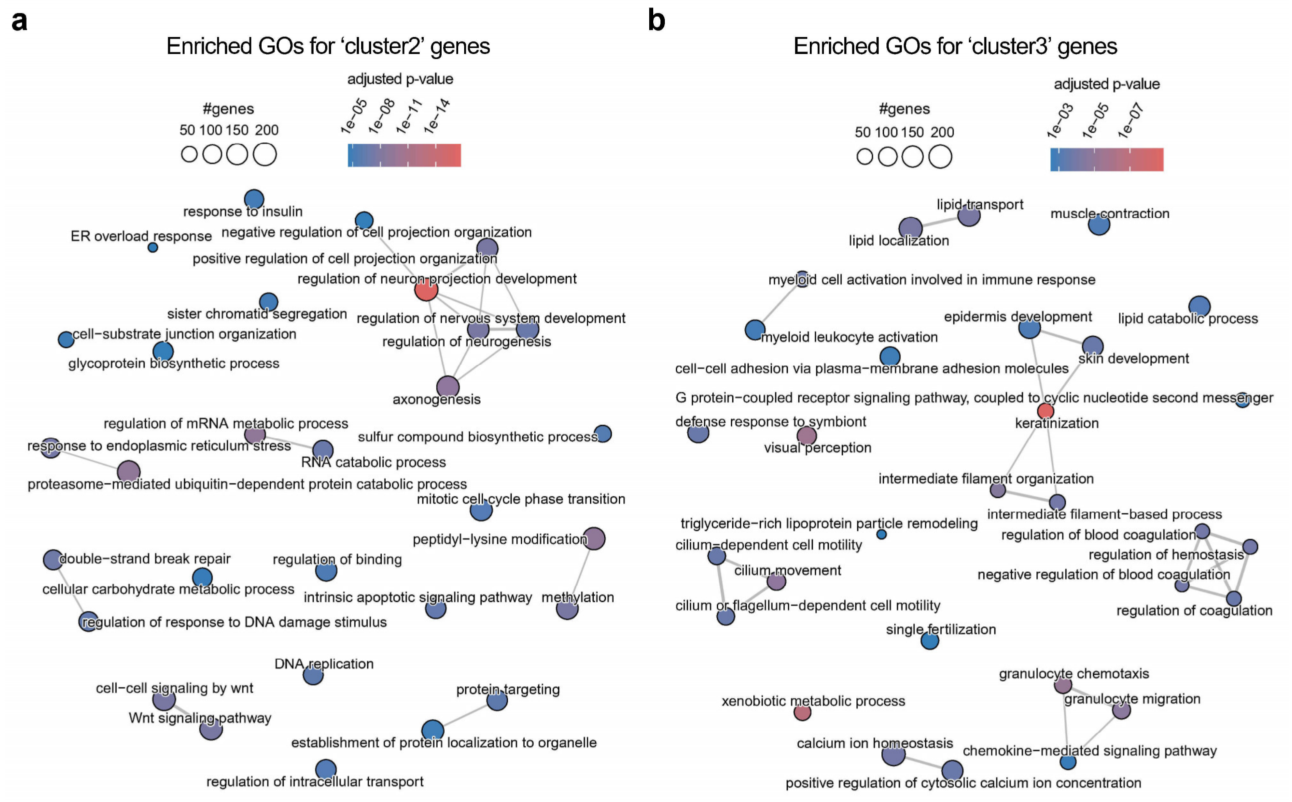


**Supplementary Figure 20 Enriched GO terms for genes of 'cluster2' and 'cluster'3 in Fig. 6b, respectively.**

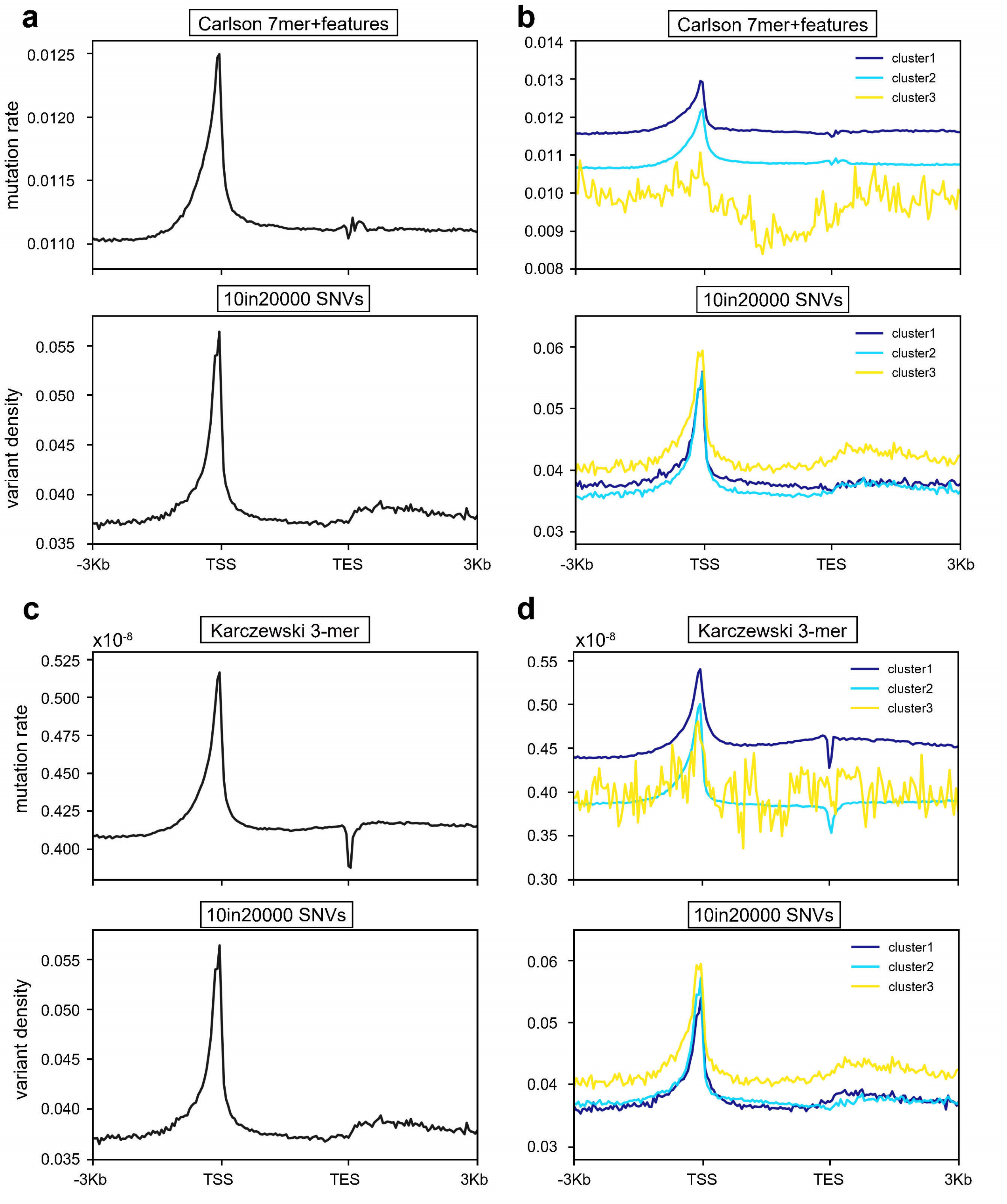


**Supplementary Figure 21 Clustering human coding genes based on mutation rate profiles of 'Carlson 7mer+features' and 'Karczewski 3-mer' models.** (**a**) Meta-gene mutation rate plots with a bin size of 50bp for regions around human coding genes. The upper is based on scaled 'Carlson 7mer+features'mutation rates, and the lower based on '10in20000' rare variants. TSS, transcription start site; TES, transcription end site. (**b**) Meta-gene plots of three gene clusters after performing K-means clustering (K=3) with 'Carlson 7mer+features' mutation rates (same data in panel **a**). The same clusters were used for generating meta-gene plots for '10in20000' data (lower panel). It is clear that the orders of clusters in the upper and lower panels are rather different. For instance, 'cluster3' genes have lowest mutation rates in the upper panel but highest rates in the lower panel, suggesting inaccuracies in ‘Carlson 7mer+features’ data. (**c-d**) Similar to panels **a-b**, except that ‘Karczewski 3-mer’ mutation rates were used for analysis.

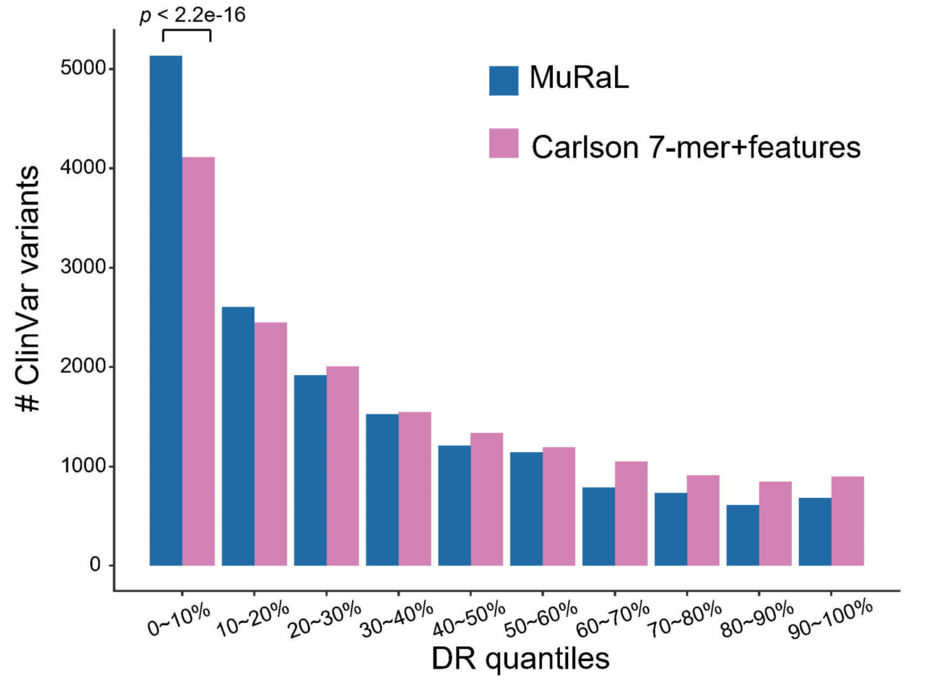


**Supplementary Figure 22 Numbers of ClinVar variants ('Pathogenic' and 'Likely pathogenic' in annotations) in regions of different DR scores.** DR scores were calculated using MuRaL and ‘Carlson 7mer+features’ mutation rates, respectively. Two-sided Fisher’s exact test was used for comparing the numbers of variants in regions with lowest 10% DR scores.

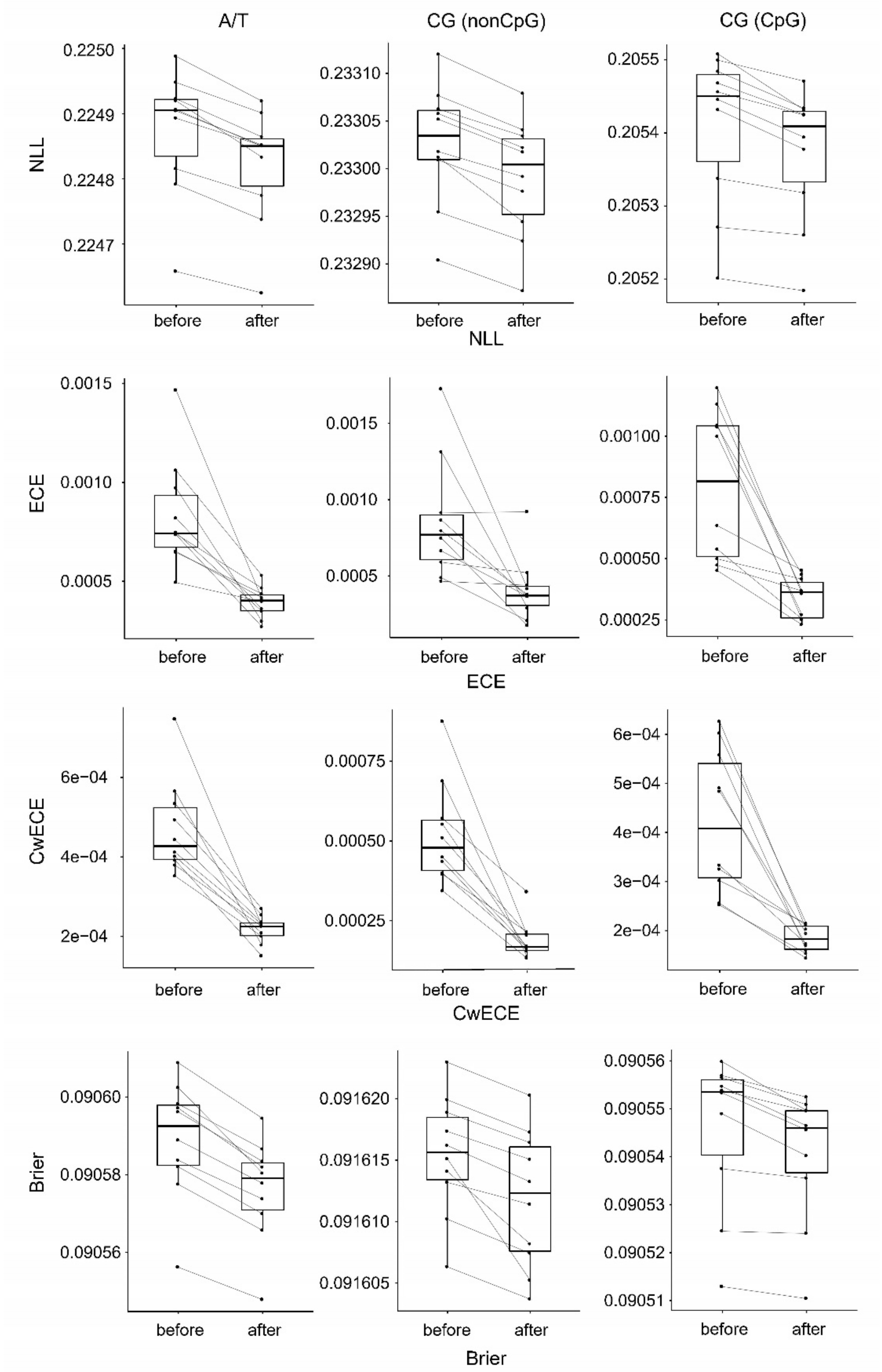


**Supplementary Figure 23 Comparison of multiple calibration-related metrics before and after applying Dirichlet calibration to the predicted mutation probabilities of validation data by the full MuRaL models**. Separate Dirichlet calibrators were estimated for models of AT, non-CpG and CpG models, respectively. ECE is short for Expected Calibration Error. Paired lines indicate values from same trials.

**Supplementary Table 1 Basic statistics for different rare variant datasets and DNMs**. Variants with too low (<15) or too high (>45) gnomAD read coverage were excluded.

| | A/T | | | | non-CpG | | | | CpG | | | |
|---|---|---|---|---|---|---|---|---|---|---|---|---|
| | **A>C** | **A>G** | **A>T** | **total** | **C>A** | **C>G** | **C>T** | **total** | **C>A** | **C>G** | **C>T** | **total** |
| 1in200 | 512,548 | 1,930,541 | 489,730 | 2,932,819 | 607,011 | 580,168 | 1,785,054 | 2,972,233 | 73,826 | 59,329 | 1,220,738 | 1,353,893 |
| 1in2000 | 1,525,079 | 5,661,078 | 1,435,767 | 8,621,924 | 1,830,683 | 1,746,677 | 5,154,321 | 8,731,681 | 210,667 | 167,941 | 3,282,065 | 3,660,673 |
| 1in7000 | 3,292,911 | 12,051,439 | 3,076,416 | 18,420,766 | 3,942,586 | 3,764,023 | 10,958,577 | 18,665,186 | 438,830 | 348,205 | 5,868,297 | 6,655,332 |
| 5in1000 | 1,589,954 | 5,986,397 | 1,515,639 | 9,091,990 | 1,897,649 | 1,816,388 | 5,508,912 | 9,222,949 | 221,580 | 177,742 | 3,736,460 | 4,135,782 |
| 10in20000 | 8,920,744 | 32,802,000 | 8,321,557 | 50,044,301 | 10,615,721 | 10,144,107 | 29,765,368 | 50,525,196 | 1,051,520 | 840,666 | 15,517,079 | 17,409,265 |
| DNMs | 30,226 | 115,404 | 28,750 | 174,380 | 36,142 | 37,143 | 105,350 | 178,635 | 3,776 | 3,298 | 76,193 | 83,267 |

**Supplementary Table 2 Training and validation mutations for human MuRaL models**. '1in2000' rare variant data were for training human AT and non-CpG models and the '5in1000' data for training the CpG model.

| | A/T (1in2000) | | | | non-CpG (1in2000) | | | | CpG (5in1000) | | | |
|---|---|---|---|---|---|---|---|---|---|---|---|---|
| | **A>C** | **A>G** | **A>T** | **total** | **C>A** | **C>G** | **C>T** | **total** | **C>A** | **C>G** | **C>T** | **total** |
| training | 88,818 | 328,381 | 82,801 | 500,000 | 105,057 | 100,209 | 294,734 | 500,000 | 26,848 | 21,275 | 451,877 | 500,000 |
| validation | 8,964 | 32,711 | 8,325 | 50,000 | 10,503 | 9,913 | 29,584 | 50,000 | 2,646 | 2,163 | 45,191 | 50,000 |

**Supplementary Table 3 Training data of different models for inferring human mutation rate maps.**

| **Model** | **Sequence space** | **Require functional genomic data?** | **# training variants** |
|---|---|---|---|
| MuRaL | ±1000 + 1 = 2001 bp | no | 1.5 millions |
| Carlson 7-mer+features | ±3 + 1 = 7 bp | yes | ~35 millions |
| Carlson 7-mer | ±3 + 1 = 7 bp | no | ~35 millions |
| Aggarwala 7-mer | ±3 + 1 = 7 bp | no | 6~11 million |
| Karczewski 3-mer | ±1 +1 = 3 bp | yes | ~24 millions |

**Supplementary Table 4 Training and validation mutations for transfer learning models with human DNMs**.

| | A/T (DNMs) | | | | non-CpG (DNMs) | | | | CpG (DNMs) | | | |
|---|---|---|---|---|---|---|---|---|---|---|---|---|
| | **A>C** | **A>G** | **A>T** | **total** | **C>A** | **C>G** | **C>T** | **total** | **C>A** | **C>G** | **C>T** | **total** |
| training | 25,911 | 99,496 | 24,593 | 150,000 | 30,364 | 31,217 | 88,419 | 150,000 | 2,237 | 1,947 | 45,816 | 50,000 |
| validation | 2,592 | 10,011 | 2,397 | 15,000 | 3,047 | 3,074 | 8,879 | 15,000 | 652 | 580 | 13,768 | 15,000 |

**Supplementary Table 5 Rare variant data of *M. mulatta* used in analysis.**

| | A/T | | | | non-CpG | | | | CpG | | | |
|---|---|---|---|---|---|---|---|---|---|---|---|---|
| | **A>C** | **A>G** | **A>T** | **total** | **C>A** | **C>G** | **C>T** | **total** | **C>A** | **C>G** | **C>T** | **total** |
| all | 1,469,900 | 4,805,114 | 1,379,786 | 7,654,800 | 1,696,924 | 1,420,709 | 5,825,832 | 8,943,465 | 276,133 | 256,390 | 5,814,742 | 6,347,265 |
| training | 96,067 | 314,061 | 89,872 | 500,000 | 95,022 | 79,317 | 325,661 | 500,000 | 21,661 | 20,138 | 458,201 | 500,000 |

| | | | | | | | | | | | | |
|---|---|---|---|---|---|---|---|---|---|---|---|---|
| validation | 9,664 | 31,298 | 9,038 | 50,000 | 9,424 | 7,935 | 32,641 | 50,000 | 2,220 | 1,983 | 45,797 | 50,000 |
| transfer learning | 28,870 | 94,241 | 26,889 | 150,000 | 28,395 | 23,680 | 97,925 | 150,000 | 6,534 | 6,067 | 137,399 | 150,000 |

**Supplementary Table 6 Rare variant data of *Arabidopsis thaliana* used in analysis.**

| | A/T | | | | non-CpG | | | | CpG | | | |
|---|---|---|---|---|---|---|---|---|---|---|---|---|
| | A>C | A>G | A>T | total | C>A | C>G | C>T | total | C>A | C>G | C>T | total |
| all | 385,740 | 728,890 | 561,161 | 1,675,791 | 569,348 | 417,419 | 1,640,805 | 2,627,572 | 88,926 | 67,096 | 384,555 | 540,577 |
| training | 22,941 | 43,614 | 33,445 | 100,000 | 21,679 | 15,777 | 62,544 | 100,000 | 16,228 | 12,408 | 71,364 | 100,000 |
| validation | 2,275 | 4,354 | 3,371 | 10,000 | 2,265 | 1,584 | 6,151 | 10,000 | 1,655 | 1,227 | 7,118 | 10,000 |

**Supplementary Table 7 Rare variant data of *D. melanogaster* used in analysis.**

| | A/T | | | | C/G | | | |
|---|---|---|---|---|---|---|---|---|
| | A>C | A>G | A>T | total | C>A | C>G | C>T | total |
| all | 51,754 | 118,919 | 114,701 | 285,374 | 119,677 | 64,684 | 234,352 | 418,713 |
| training | 18,009 | 41,661 | 40,330 | 100,000 | 28,414 | 15,660 | 55,926 | 100,000 |
| validation | 1,857 | 4,069 | 4,074 | 10,000 | 2,863 | 1,572 | 5,565 | 10,000 |

**Supplementary Table 8 Configuration of key hyperparameters for training human MuRaL models.** Same hyperparameter settings were used for AT, non-CpG and CpG models. 'tune.loguniform' is a function in Ray Tune for sampling values for corresponding hyperparameters. The transfer learning models were trained with human DNMs.

| hyperparameter | *ab initio* models | transfer learning models |
|---|---|---|
| local_radius | 7 | 7 |
| local_order | 3 | 3 |
| local_hidden1_size | 150 | 150 |
| local_hidden2_size | 75 | 75 |
| distal_radius | 1000 | 1000 |
| CNN_kernel_size | 3 | 3 |
| CNN_out_channels | 32 | 32 |
| learning_rate | 0.001 | tune.loguniform(1e-5, 1e-3) |
| weight_decay_auto | 0.1 | 0.1 |
| LR_gamma | 0.95 | 0.95 |

**Supplementary Table 9 Configuration of key hyperparameters for training MuRaL models of *M. mulatta*.** Same hyperparameter settings were used for AT, non-CpG and CpG models. 'tune.loguniform' is a function in Ray Tune for sampling values for corresponding hyperparameters. The transfer learning models were trained with monkey rare variants and human MuRaL models.

| hyperparameter | *ab initio* models | transfer learning models |
|---:|---:|---:|
| local_radius | 7 | 7 |
| local_order | 3 | 3 |
| local_hidden1_size | 150 | 150 |
| local_hidden2_size | 75 | 75 |
| distal_radius | 1000 | 1000 |
| CNN_kernel_size | 3 | 3 |
| CNN_out_channels | 32 | 32 |
| learning_rate | 0.001 | tune.loguniform(1e-5, 1e-3) |
| weight_decay_auto | 0.1 | 0.1 |
| LR_gamma | 0.95 | 0.95 |

**Supplementary Table 10 Configuration of key hyperparameters for training MuRaL models of *A. thaliana.*** Same hyperparameter settings were used for AT, non-CpG and CpG models.

| hyperparameter | AT model / non-CpG model / CpG model |
|---:|---:|
| local_radius | 7 |
| local_order | 3 |
| local_hidden1_size | 150 |
| local_hidden2_size | 75 |
| distal_radius | 1000 |
| CNN_kernel_size | 3 |
| CNN_out_channels | 32 |
| learning_rate | 0.001 |
| weight_decay_auto | 0.1 |
| LR_gamma | 0.95 |

**Supplementary Table 11 Configuration of key hyperparameters for training MuRaL models of *D. melanogaster.*** Same hyperparameter settings were used for AT and CG models.

| hyperparameter | AT model / CG model |
|---:|---:|
| local_radius | 7 |
| local_order | 3 |
| local_hidden1_size | 150 |
| local_hidden2_size | 75 |
| distal_radius | 1000 |
| CNN_kernel_size | 3 |
| CNN_out_channels | 32 |
| learning_rate | 0.001 |
| weight_decay_auto | 0.1 |
| LR_gamma | 0.95 |
